# Overriding Mendelian inheritance in *Arabidopsis* with a CRISPR toxin-antidote gene drive that impairs pollen germination

**DOI:** 10.1101/2023.10.10.561637

**Authors:** Yang Liu, Bingke Jiao, Jackson Champer, Wenfeng Qian

**Author notes:** Correspondence to: Wenfeng Qian Institute of Genetics and Developmental Biology, Chinese Academy of Sciences Beijing 100101, China.

## Abstract

Synthetic gene drives, inspired by natural selfish genetic elements, present transformative potential for disseminating traits that benefit humans throughout wild populations, irrespective of potential fitness costs. Here, we constructed a gene drive system called CRISPR-Assisted Inheritance utilizing *NPG1* (*CAIN*), which employs a toxin-antidote mechanism in the male germline to override Mendelian inheritance in plants. Specifically, a gRNA-Cas9 cassette targets the essential *No Pollen Germination 1* (*NPG1*) gene, serving as the toxin to block pollen germination. A recoded, CRISPR-resistant copy of *NPG1* serves as the antidote, providing rescue only in pollen cells that carry the drive. To limit potential consequences of inadvertent release, we used self-pollinating *Arabidopsis thaliana* as a model. The drive demonstrated a robust 88–99% transmission rate over two successive generations, producing minimal resistance alleles that are unlikely to inhibit drive spread. Our study provides a strong basis for rapid genetic modification or suppression of outcrossing plant populations.

## Introduction

Facing diverse challenges such as disease control, food security threats from agricultural pests or weeds, and the environmental crisis of lost biodiversity, the genetic manipulation of wild populations has emerged as potentially powerful and transformative strategy. However, efforts to introduce traits that benefit humans are almost invariably limited by classical Mendelian inheritance and Darwinian selection because they are selectively neutral or even detrimental to the organisms themselves and are therefore rapidly lost in targeted populations. Nevertheless, selfish genetic elements are prevalent in nature and can be transmitted to progeny at super-Mendelian (>50%) frequencies^1-3^. Inspired by these natural processes, synthetic gene drives have been proposed for spreading genetic alterations throughout populations in their natural environment, despite potential fitness costs^4-7^. Thus, gene drive technologies offer a compelling tool to address several global challenges.

Synthetic gene drive systems have been implemented for population modification or suppression in various hosts, including yeast^8,9^, mosquitoes^10-15^, flies^16-18^, and mice^19,20^. Among these, most are homing-based drives and have been shown to spread rapidly within populations^8,10-16,19^. This propagation is predominantly driven by the “copy-and-paste” mechanism that depends on CRISPR-Cas9 mediated DNA cleavage (*i.e.*, double strand break, DSB) of the homologous allele followed by repair via homology-directed recombination (HDR), thus converting diploid germline cells from heterozygotes to homozygotes. Nevertheless, homing-based drives frequently yield resistance alleles, defined as those which, due to the activity of endogenous end-joining DNA repair pathways, become resistant to further cleavage. Notably, this repair pathway is preferentially used in plants^21^, thus posing a significant challenge to sustained dissemination of homing-based gene drives in plant populations.

To develop an efficient synthetic gene drive for plants, it is therefore necessary to circumvent dependency on the HDR repair pathway. One solution is the utilization of toxin-antidote gene drives, exemplified by naturally occurring systems such as the *t*-haplotype in mice^22,23^. These gene drives typically feature a toxin, expressed prior to meiosis and consequently disseminated to all gametes, disrupting regular gametogenesis. The genetically linked antidote is activated in a later, post-meiotic stage to mitigate the toxin-induced damage, conferring a fitness advantage to its carrier. Although natural toxin-antidote gene drives involve sophisticated regulatory dynamics that might not seamlessly transfer across species, CRISPR-Cas9 could overcome this obstacle and provide a universal template for mimicking toxin-antidote strategies. In such a design, a CRISPR-Cas9 cassette could act as a toxin, cleaving an essential gene to produce a loss-of-function allele, while a recoded, Cas9-resistant version of the essential gene functions as the antidote to rescue the defective functionality^17,18,24^.

Examples of this strategy are *Cleave and Rescue* (*ClvR*)^17^ and Toxin-Antidote Recessive Embryo (TARE)^18,24^. These drives target a haplosufficient gene essential for zygotic development, resulting in the demise of progeny lacking the drive but carrying two loss-of-function alleles of the essential gene. In contrast, all progeny inheriting the drive allele survive. Despite having several advantages, the efficiency of these drives is limited by the frequency-dependent mechanism, necessitating higher release frequencies and rendering them inappropriate for strong suppression^24^. The hypothetical Toxin-Antidote Dominant Sperm (TADS) drive, designed to disrupt genes essential in spermatogenesis, provides a theoretical basis for improving the power of toxin-antidote systems^24^. This design enables rapid drive propagation and even potential suppression if the drive locates in a haplosufficient gene essential for male fertility (where drive homozygotes would render male sterile). However, this approach has never been experimentally demonstrated, most likely due to the inherent challenges of identifying target genes with haploid stage expression and exclusively required for sperm development.

In this work, we present *CAIN* (CRISPR-Assisted Inheritance utilizing *NPG1*), a synthetic toxin-antidote gene drive developed for plants. The *CAIN* design exploits the prolonged haploid stage in plants to target an essential gene in pollen germination, *No Pollen Germination 1* (*NPG1*). We introduced *CAIN* into *A. thaliana* and monitored its transmission through a red fluorescent seed phenotype, encoded within the drive. *CAIN* transmission rates greatly exceed the expected Mendelian inheritance of 50% in heterozygotes, reaching 88-99% within two successive generations. Genotyping revealed a scarcity of resistance alleles, and modeling predicts the drive could rise from 1% to 99% prevalence within roughly 17 generations. The rapid spread of *CAIN* in *A. thaliana* demonstrates its potential for broad applications in diverse plant species, paving the way for innovative approaches in ecological management and sustainable agriculture.

## Results

### Design of *CAIN*, a CRISPR-based toxin-antidote gene drive targeting pollen germination

*CAIN* consists of two tightly linked components: the toxin and the antidote, which can optionally be combined with a cargo. The toxin is a gRNA-Cas9 cassette that can introduce loss-of-function mutations in a gene essential for pollen germination, *No Pollen Germination* (*NPG1*)^25^, while the antidote is a recoded, CRISPR-resistant version of *NPG1* (Fig. 1a, Fig. S1). To mitigate potential negative effects on plant development and fitness caused by Cas9 activity in somatic tissues, we selected two promoters with distinct temporal expression patterns for male germline expression of *Cas9* (Fig. 1a): the *DMC1* (*Disruption of Meiotic Control 1*) promoter, which expresses *Cas9* in the pollen mother cells within anthers^26^, and the *TPD1* (*Tapetum Determinant 1*) promoter, which activates gene expression in sporogenous cells that eventually develop into pollen mother cells^27^. To guarantee the effectiveness of *CAIN*’s toxicity, we incorporated four gRNAs (gRNA2, gRNA6, gRNA11, and gRNA23) of varying efficiencies into *CAIN* (Fig. S1; Table S1), each expressed under the control of a U6 promoter^28^. As the cargo, we chose the red fluorescence protein marker, FAST^29^ (*i.e.*, fluorescence-accumulating seed technology), which generates a red fluorescent signal in dry seed, to facilitate phenotypic tracking of the gene drive (Fig. 1a). Furthermore, we generated a construct containing only the FAST marker to serve as a negative control (Fig. S2).

**Fig. 1.**
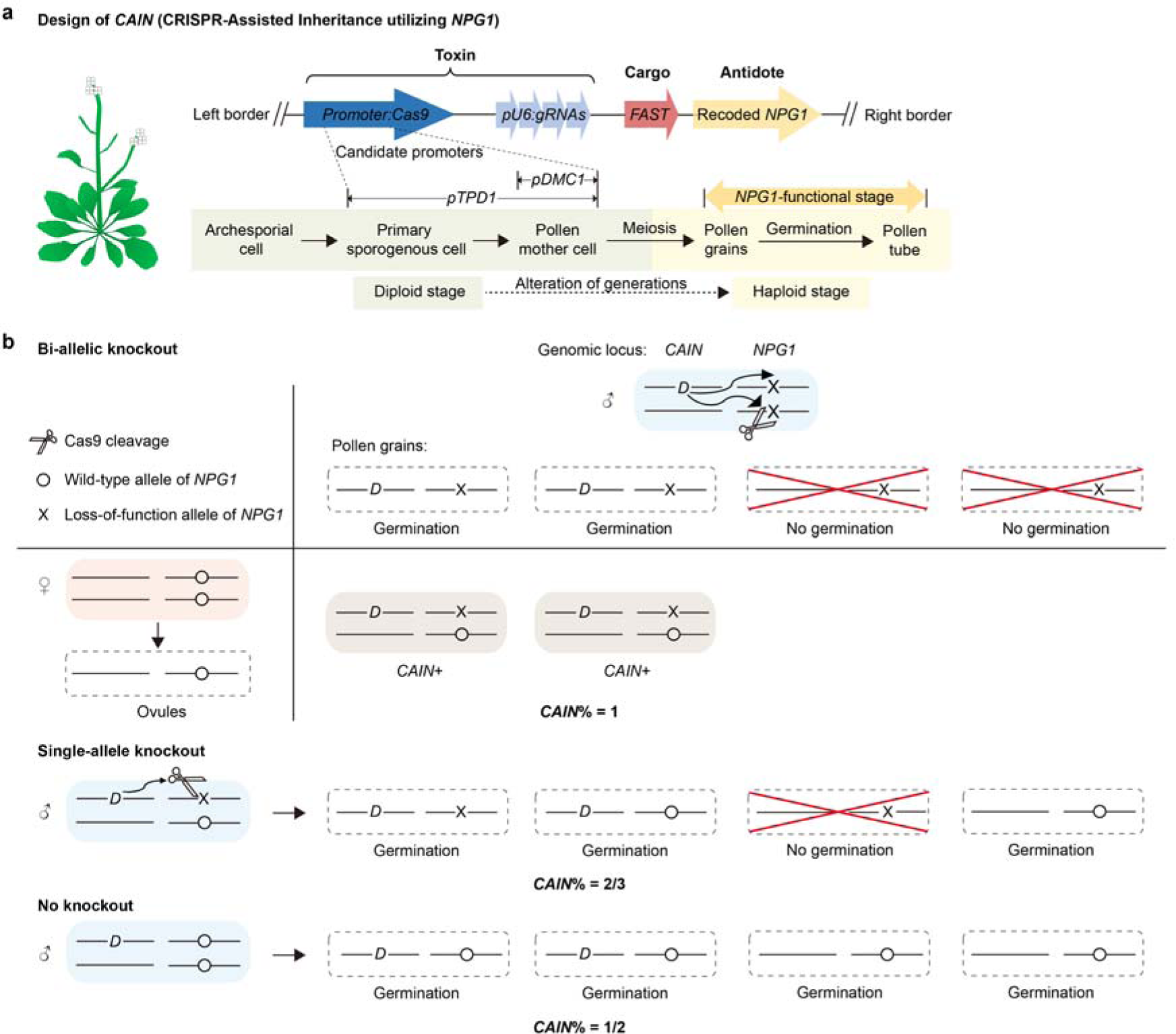
The *CAIN* gene drive design and predicted genetic behavior. **a,** The T-DNA construct of *CAIN*. The corresponding stages of *A. thaliana* male gametophyte development are depicted below. The *TPD1* promoter provides a longer window for the toxin to function in progenitor pollen cells, potentially facilitating higher overall Cas9 cleavage activity before *NPG1* function is first required (*i.e.*, in pollen germination). *TPD1*: *Tapetum Determinant 1*. *DMC1*: *Disruption of Meiotic Control 1*. *NPG1*: *No Pollen Germination 1*. **b,** Anticipated fractions of F1 progeny carrying *CAIN* (*CAIN*%) from a cross between a *CAIN*-carrying male parent (*CAIN*/+) and a wild-type female parent (+/+), based on various scenarios of *NPG1* cleavage in the male germline. Salmon, sky blue, and grey boxes represent female parents, male parents, and progeny, respectively. Dashed boxes depict gametophytes. Red crosses represent pollen grains that cannot germinate.

In theory, the toxin can potentially disrupt both alleles of *NPG1* prior to meiosis and therefore impair germination of all four pollen grains, and the antidote can neutralize this effect only when present within a pollen grain. As a result, when a plant carrying a single copy of *CAIN* pollinates a wild-type plant, only the two *CAIN*-carrying pollen grains can successfully complete germination, yielding 100% transmission of *CAIN* (Fig. 1b). Even if only one of the two alleles of the essential gene is disrupted by the toxin, the transmission rate of *CAIN* will be two thirds (Fig. 1b), still well above 50% Mendelian inheritance.

This process can propagate in successive generations, ultimately disseminating *CAIN* throughout the population through continuous cross-pollination. Although gene drive systems are primarily designed to function in outcrossing populations, we selected *A. thaliana*, which primarily undergoes self-pollination, as our experimental model. Throughout the entire experiment, plants carrying *CAIN* were manually crossed with wild-type plants. Coupled with rigorous experimental procedures and management, the choice of this predominantly self-pollinating species further bolsters the biosafety of our gene drive.

### Substantially increased transmission rate of *TPD-CAIN* to F1 progeny

The *CAIN* design does not require linkage between the drive and the target gene, and consequently, we used *Agrobacterium*-mediated floral dipping to introduce the *CAIN* construct (*DMC-CAIN* and *TPD-CAIN*) into wild-type *A. thaliana* Col-0 at a random locus (Fig. 2a, Fig. S3). We screened transformants (*i.e.*, T1) with the exogenous DNA sequence inserted into only a single locus (*DMC-CAIN*/+ or *TPD-CAIN*/+, where “+” represents the other allele that does not carry the drive) for subsequent characterization (Table S2), since these transformants will exhibit a 50% transmission rate according to Mendelian inheritance in the absence of gene drive activity.

**Fig. 2.**
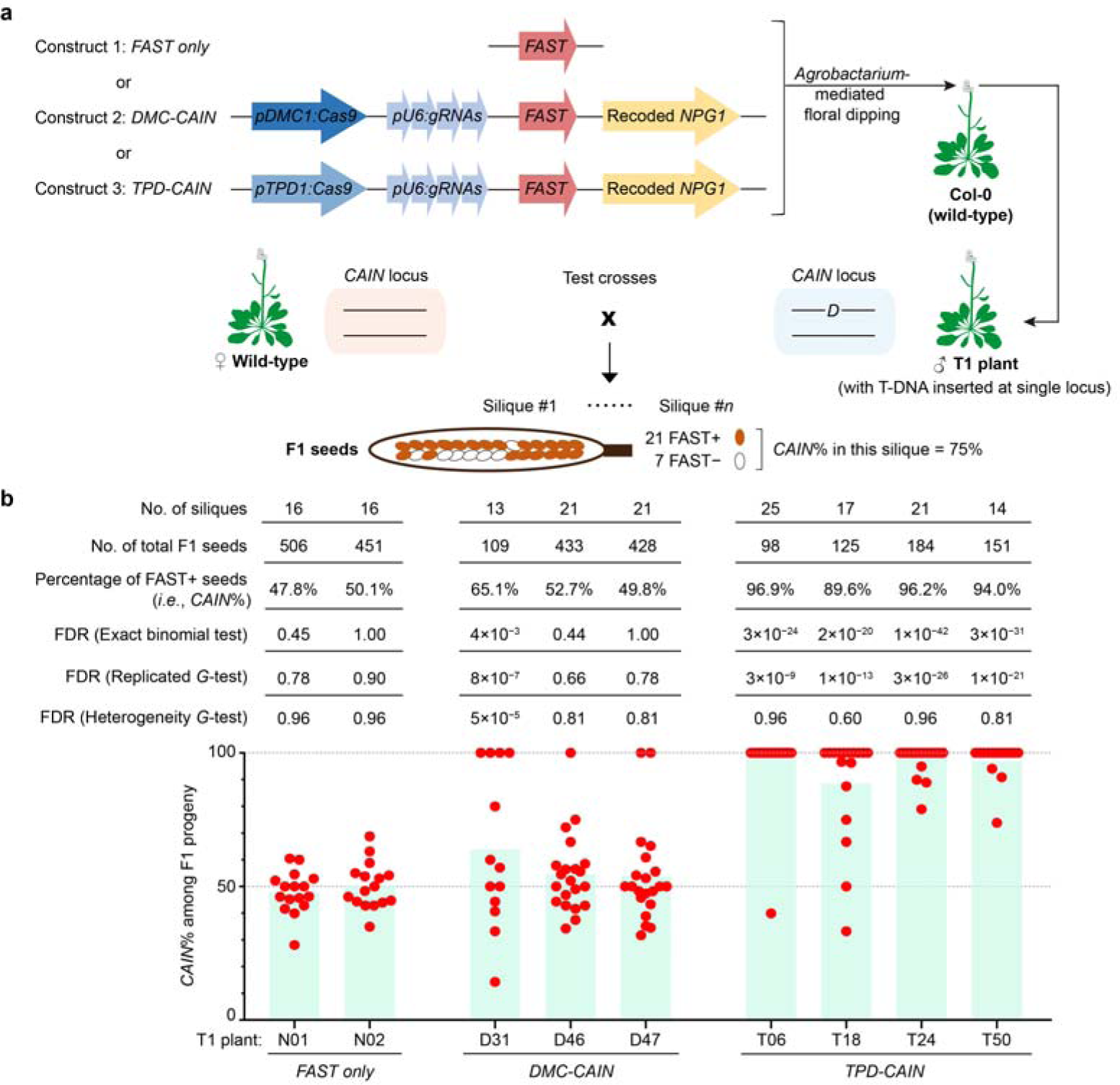
Transmission rate of *CAIN* from the T1 to F1 generation. **a,** Experimental procedure for transformation yielding T1 plants and subsequent test crosses generating F1 progeny. The transmission rate of *CAIN* (*CAIN*%) was calculated from the fraction of F1 seeds exhibiting FAST+. **b,** Bar plots show the average transmission rate for each pair of parent plants, and individual red dots indicate the transmission rate estimated from a single silique. The null hypothesis for the statistic tests was 50% transmission rate predicted by Mendelian inheritance. FDR, False discovery rate.

To evaluate whether *CAIN* can successfully enhance the likelihood of inheritance beyond Mendelian expectation through paternal transmission, we used *DMC-CAIN*/+ or *TPD-CAIN*/+ T1 plants as the paternal donor in crosses with wild-type female parent (Fig. 2a). This experiment was effectively a test cross because the female parent would always supply a wild-type allele to the F1 progeny, and therefore *CAIN* transmission from the male parent can be determined based on the dominant FAST (*i.e.*, red fluorescence in F1 seeds) phenotype. In the F1 progeny of *DMC-CAIN*/+ T1 plants, the fraction of individuals inheriting *DMC-CAIN* (*i.e.*, *CAIN*%) was significantly higher than predicted Mendelian outcomes in only one (*i.e.*, D31, *CAIN*% = 65.1%, false discovery rate = 0.004, binomial test) of the three crosses (Fig. 2b; Table S3). In comparison, 89.6% to 96.9% of F1 progeny in all four crosses of the *TPD-CAIN*/+ T1 plants inherited *TPD-CAIN*, indicating a substantial deviation from Mendelian inheritance (Fig. 2b; Table S3). By contrast, the negative control construct (*i.e.*, FAST only) was indeed transmitted at a Mendelian rate of approximately 50% (Fig. 2b; Table S3).

Because transmission rates in each silique can be treated as the results of an independent test, we further determined the significant deviation from 50% transmission rate using replicated tests of goodness of fit (*G*-test, Fig. 2b) to account for variation in transmission rates among siliques. We found that the transmission rate of *DMC-CAIN* was significantly heterogeneous among siliques of male parent D31 (*P* = 5 × 10^−5^, heterogeneity *G*-test, Fig. 2b), which was the only cross showing >50% transmission rate. This heterogeneity is likely due to high stochasticity in Cas9 cleavage efficiency, leading to non-trivial variability in *NPG1* cleavage status between pollen from different anthers or flowers. In contrast, no significant heterogeneity in transmission rate was detected among siliques of male parent *TPD-CAIN*/+ T1 plants. Moreover, the F1 populations of all four crosses showed *CAIN*% close to 100% (Fig. 2b), suggesting robust performance by *TPD-CAIN*.

### Transmission bias in *TPD-CAIN* is determined by direction of pollination

To assess whether *TPD-CAIN* biased Mendelian inheritance by disrupting *NPG1* as intended (Fig. 1), we genotyped F1 plants by Sanger sequencing of the region around the Cas9 target sites in *NPG1* in somatic tissues (*i.e.*, rosette leaves or early inflorescence, Fig. 3a). Genotyping of FAST+ (*i.e.*, *TPD-CAIN*/+) F1 plants (*n* = 49) revealed that all carried a putative *NPG1* loss-of-function allele (*NPG1*^−^), with diverse indels observed at gRNA2 and gRNA11 target sites in 96% and 100% of F1 progeny, respectively (Fig. 3b and S4; Table S4). Theoretically, indels of lengths multiples of three could create CRISPR-resistant *NPG1* alleles yet preserving the reading frame, but they were rare (6% at gRNA2 and 4% at gRNA11 target site, Table S4). This rarity, especially across both gRNA sites (*i.e.*, 0%, Table S4), indicated that the *CAIN* design likely had low rates of functional resistance allele formation, particularly with its incorporation of multiple gRNAs. The ubiquity of *NPG1*^−^ in almost every F1 plant suggested the high efficiency of CRISPR-based *NPG1* knockout, which should further impair pollen germination, the mechanism underlying biased transmission of *TPD-CAIN*.

**Fig. 3.**
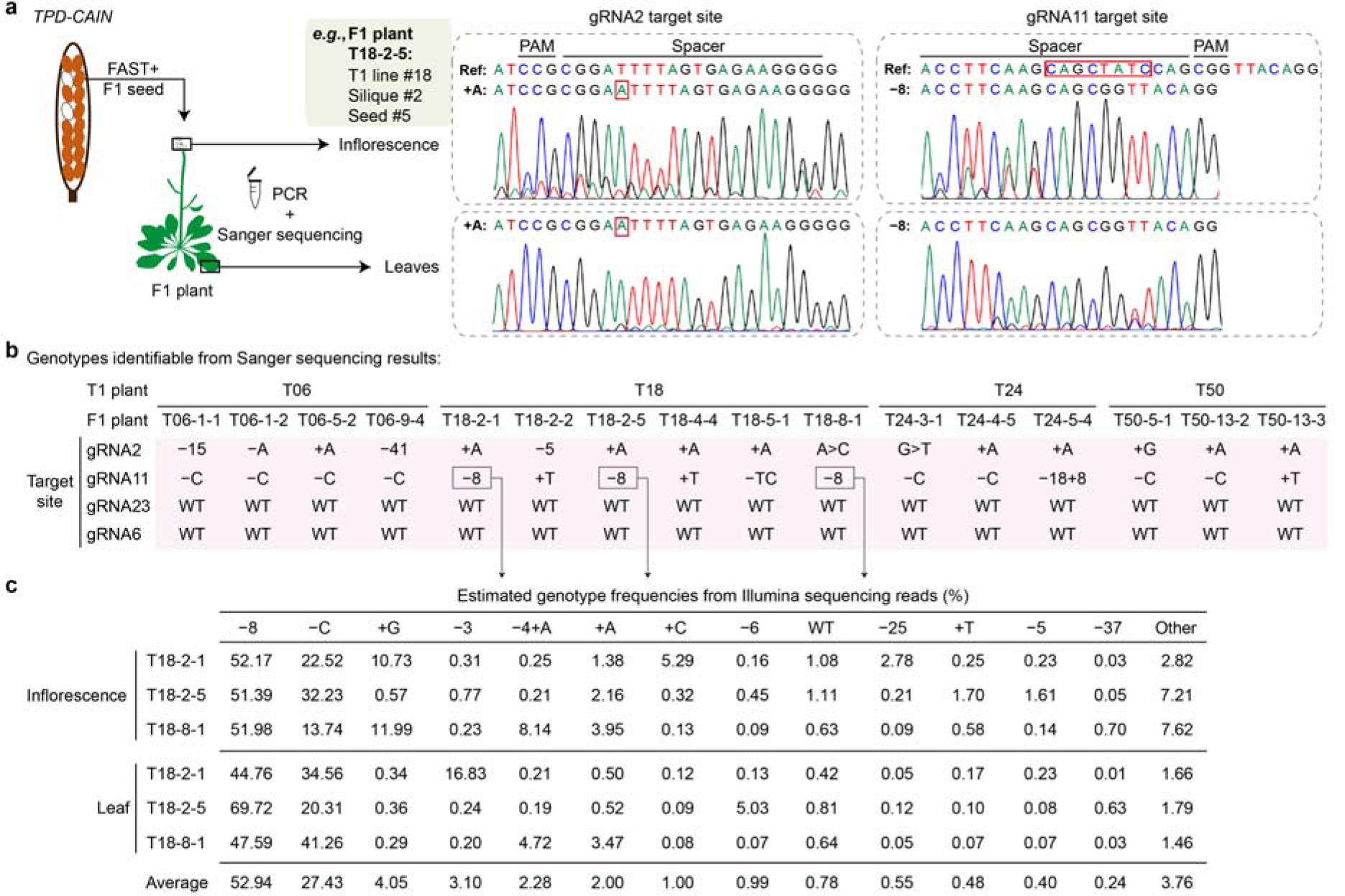
Genotypes at the *NPG1* locus for FAST+ F1 plants generated from male T1 plants carrying *TPD-CAIN*. **a,** Diagram of the somatic tissues sampled for genotyping F1 plants, with example chromatographs. **b,** Sanger sequencing-based genotyping results for 16 FAST+ F1 plants at four gRNA target sites. Symbols “+” and “−” before numerical numbers and bases denote insertion and deletion events, respectively. Numerical values indicate the number of nucleotides involved in the indels for cases where the count exceeds two. The notation “A>C” represents a base substitution from adenine (A) to cytosine (C). “WT” represents the wild-type allele. **c,** Illumina sequencing-based genotyping results for the gRNA11 target site.

Subsequent reciprocal crosses between *TPD-CAIN*/+ F1 and the wild type (+/+) were conducted to determine if the biased inheritance of *TPD-CAIN* was influenced by the direction of pollination. When *TPD-CAIN*/+ F1 plants served as male parent (*n* = 13), *TPD-CAIN* transmission rates were significantly higher than that expected in Mendelian inheritance (*i.e.*, 50%, Fig. 4a; Table S3). In contrast, in crosses with *TPD-CAIN*/+ F1 plants serving as the female parent (*n* = 8), the *TPD-CAIN* transmission rate did not significantly deviate from 50% (Fig. 4b; Table S3). These findings collectively indicated that the biased transmission of *TPD-CAIN* was likely due to defective male gametophyte resulting from *NPG1* knockout.

**Fig. 4.**
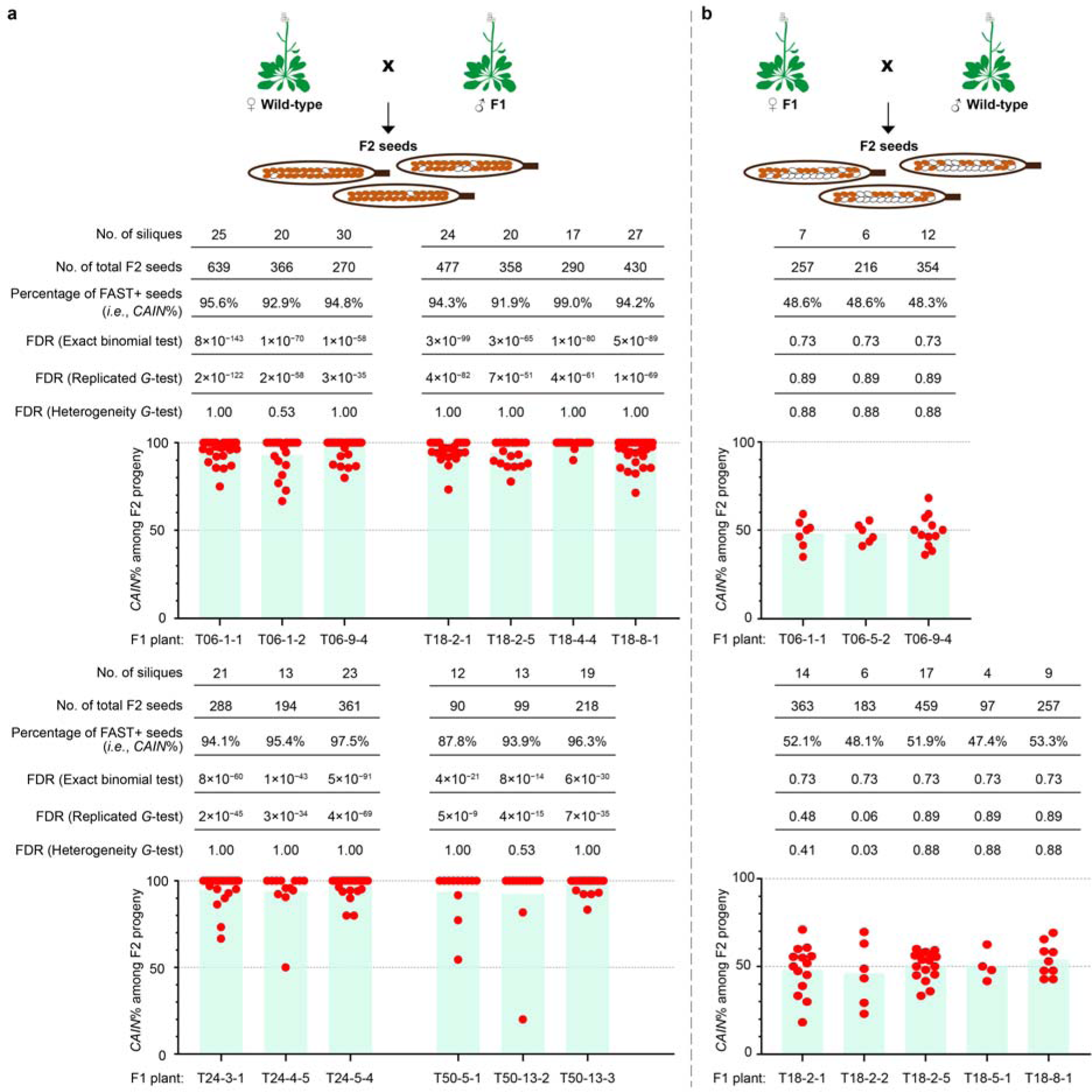
Transmission rate of *TPD-CAIN* from the F1 to F2 generation in reciprocal crosses. Average *TPD-CAIN* transmission rate depicted in bar plots with FAST+ F1 plants serving as either the (**a**) male or (**b**) female parent.

### The efficacy of *CAIN* is intrinsically tied to cleavage efficiency at the *NPG1* locus

Two potential explanations underlie the lower transmission rates of *DMC-CAIN* compared to *TPD-CAIN* (Fig. 2). First, Cas9 expression from the *DMC1* promoter may result in lower Cas9 activity than that driven by the *TPD1* promoter, and thus reduced DNA cleavage at the *NPG1* locus. Second, the timing of Cas9 activity might occur too late if expressed from the *DMC1* promoter, allowing sufficient expression of the native, undisrupted *NPG1* for the germination of some pollen grains. To differentiate between these hypotheses, we closely examined *NPG1* genotype in *DMC-CAIN*/+ F1 plants (*n* = 18, Table S4). Our analyses revealed that only two F1 plants were *NPG1*^−/−^ (D31-7-1 and D31-7-2), one plant was *NPG1*^+/−^ (D31-8-3), and the remaining 15 plants were *NPG1*^+/+^ (*NPG1*^+^ representing the wild-type allele of *NPG1*, Fig. 5a–5b; Table S4). These findings indicated that cleavage efficiencies were notably lower at the *NPG1* locus in *DMC-CAIN*/+ plants than in *TPD-CAIN*/+ plants (Fig. 3b), providing a plausible explanation for the varying transmission rates between the two *CAIN* constructs, without necessarily invoking timing-based explanation.

**Fig. 5.**
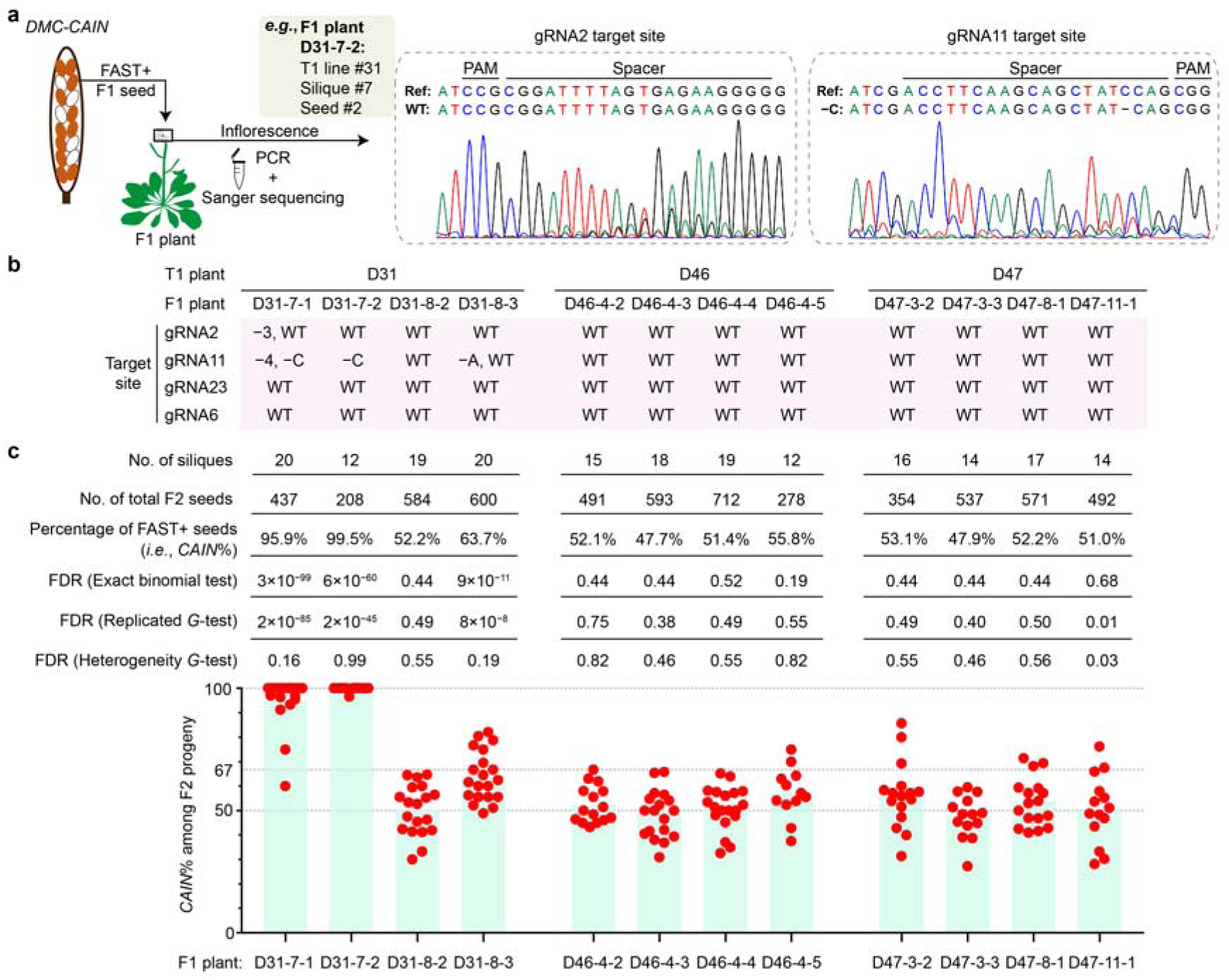
Transmission rate of *DMC-CAIN* from the F1 to F2 generation. **a,** Diagram of genotyping in F1 plants’ inflorescence. **b,** Sanger sequencing-based genotyping results for 12 FAST+ F1 plants at four gRNA target sites. **c,** Bar plots illustrating the average *DMC-CAIN* transmission rate in F2 seeds with FAST+ F1 as the male parent.

We next examined whether *DMC-CAIN*, despite its lower Cas9-mediated DNA cleavage efficiency, could still confer biased transmission across subsequent generations. For this purpose, we used two *DMC-CAIN*/+; *NPG1*^−/−^ F1 plants (D31-7-1 and D31-7-2) to pollinate wild-type plants, then screened for red fluorescence in F2 seeds of both crosses (Fig. 5b–5c; Table S3). *DMC-CAIN* transmission rates reached 95.9% and 99.5% in F2 progeny of each respective cross. In contrast, the *CAIN*% was 63.7% (approximately 2/3) in the F2 progeny generated from male parent *DMC-CAIN*/+; *NPG1*^+/−^ F1 plant D31-8-3 (Fig. 5b–5c; Table S3). F2 progeny of the remaining nine *DMC-CAIN*/+; *NPG1*^+/+^ F1 plants showed a *CAIN*% close to 50% (Fig. 5b–5c; Table S3). On average, the F1 to F2 transmission of *DMC-CAIN* is 57.5% (3367/5857, Table S3). This limited transmission rate underscored the crucial role of cleavage efficiency in determining the efficacy of the *CAIN* system.

### *TPD-CAIN* transmission rates <100% are attributable to failed DNA cleavage events and incomplete penetrance

Transmission rates of *TPD-CAIN* from male T1 parents ranged from 89.6% to 96.9% (Fig. 2b; Table S3), indicating 3.1%–10.4% of F1 progeny did not inherit *TPD-CAIN* (*i.e.*, FAST−), while 1.0%–12.2% of F2 progeny did not inherit the drive from male F1 parents (Fig. 4a; Table S3). This prompted our investigation into the causative factors resulting in failure to inherit *TPD-CAIN*. Based on the relevance of *NPG1* cleavage in *DMC-CAIN* transmission, we hypothesized that FAST− F1 plants might be the product of male gametophytes in which *NPG1* cleavage failed. We therefore scrutinized the genotypes of 11 FAST− F1 plants at the *NPG1* locus in the region around the four gRNA target sites using Sanger sequencing (Fig. 6a). Notably, six of these plants were homozygous wild type at all four sites (*i.e.*, *NPG1*^+/+^, Fig. 6b), indicating that the male T1 gametophyte contributed a *NPG1*^+^ allele. These observations strongly suggested that multiple copies of *NPG1* evaded Cas9-mediated DNA cleavage in the male gametophyte of *TPD-CAIN*/+ T1 plants, despite successful integration of the drive.

**Fig. 6.**
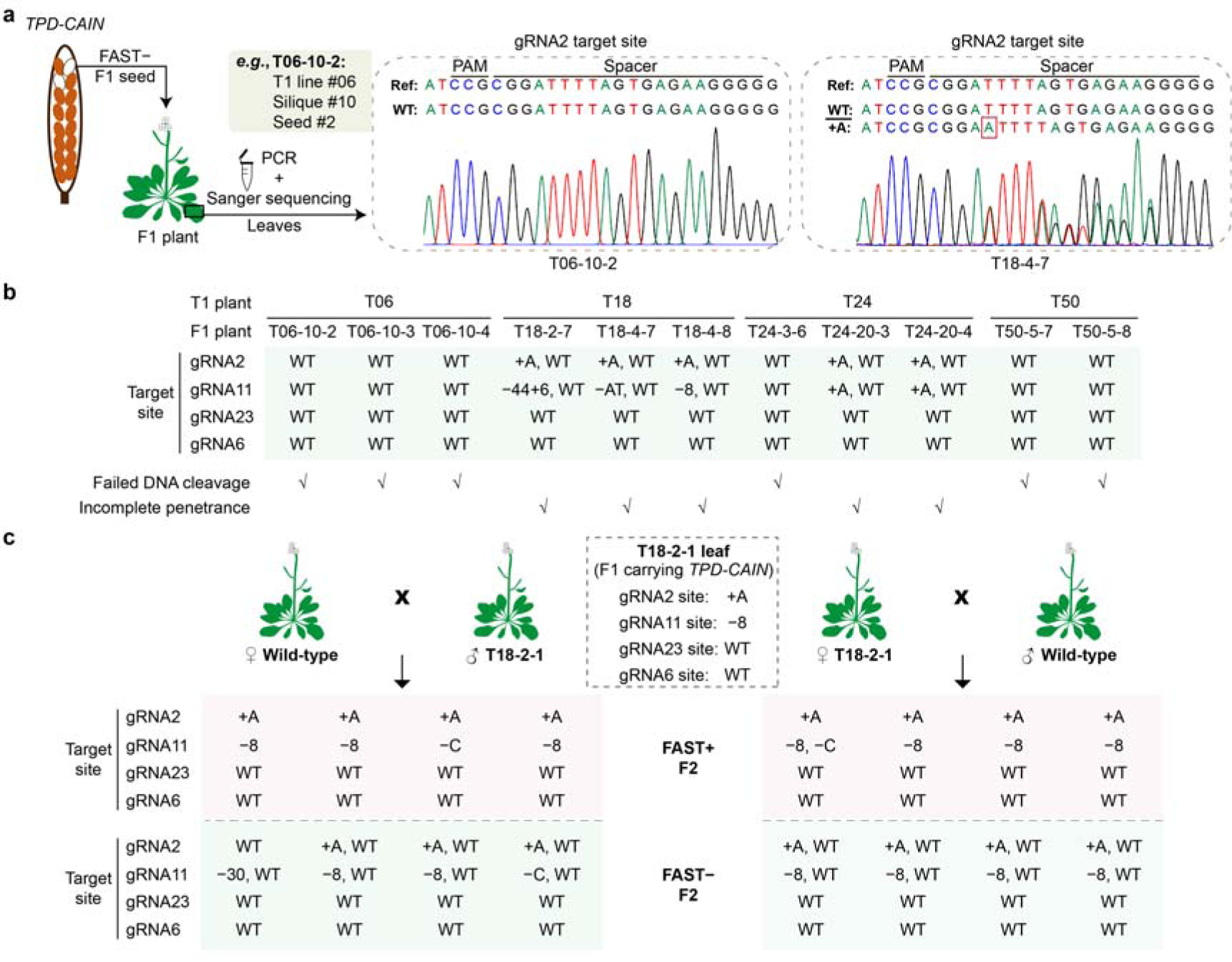
Genotypes at the *NPG1* locus for FAST− F1 and FAST+/− F2 plants. **a,** Diagram showing genotyping in the leaves of FAST− F1 plants generated from male T1 plants carrying *TPD-CAIN*. **b,** Sanger sequencing-based genotyping results for 11 FAST− F1 plants (generated from male T1 plants carrying *TPD-CAIN*) at four gRNA target sites at the *NPG1* locus. The presence of check marks (√) for each F1 plant indicates the inferred mechanisms—either failed DNA cleavage or incomplete penetrance. **c,** Sanger sequencing-based genotypes at four gRNA target sites for FAST+ and FAST− F2 plants resulting from reciprocal crosses between F1 plants carrying *TPD-CAIN* (*TPD-CAIN*/+) and the wild type (+/+).

On the other hand, five of FAST− F1 plants had a “*NPG1*^+/−^” genotype at both gRNA2 and gRNA11 sites (Fig. 6b), including three F1 progeny of T18 and two F1 progeny of T24. Considering that the female parent Col-0 can only contribute a *NPG1*^+^ allele, the male parent should have provided the *NPG1*^−^ allele through pollen transmission. It is noteworthy that post-zygotic cleavage is an unlikely explanation for the presence of these *NPG1*^−^ alleles, as *TPD-CAIN* was absent in these FAST− F1 plants. This finding suggested that some pollen grains carrying *NPG1*^−^ alleles but lacking the antidote could still germinate, indicating an incomplete penetrance of the *NPG1*^−^ sterility phenotype. This could potentially result from a lack of Cas9 cleavage in some pollen mother cells, allowing sufficient *NPG1* expression for pollen grain germination before the wild-type *NPG1* was subsequently disrupted by Cas9 cleavage.

To understand the possible population-level impacts of failed DNA cleavage and incomplete penetrance on the dynamics of *CAIN* drive dissemination, we conducted computational simulations built on a Wright-Fisher model. This individual-based, stochastic model assumed a finite, randomly mating population reproducing in discrete, non-overlapping generations. Starting with 9900 wild-type individuals and 100 *TPD-CAIN*/+ ones, we preserved population size, randomly chose mating pairs, and confined DNA cleavage and transmission bias to male parents. Taking into account the factors of male germline cleavage efficiency and penetrance rate estimated for *TPD-CAIN* (98.4% and 96.0%, respectively; Fig. S5a), the simulation results indicated that its spread from 1% to 99% of the population would take approximately 17 generations, only one generation longer than for optimal conditions (*i.e.*, 100% DNA cleavage efficiency and complete penetrance, Fig. 7a), indicating that the efficiency of *TPD-CAIN* spread remained relatively robust despite these influences.

**Fig. 7.**
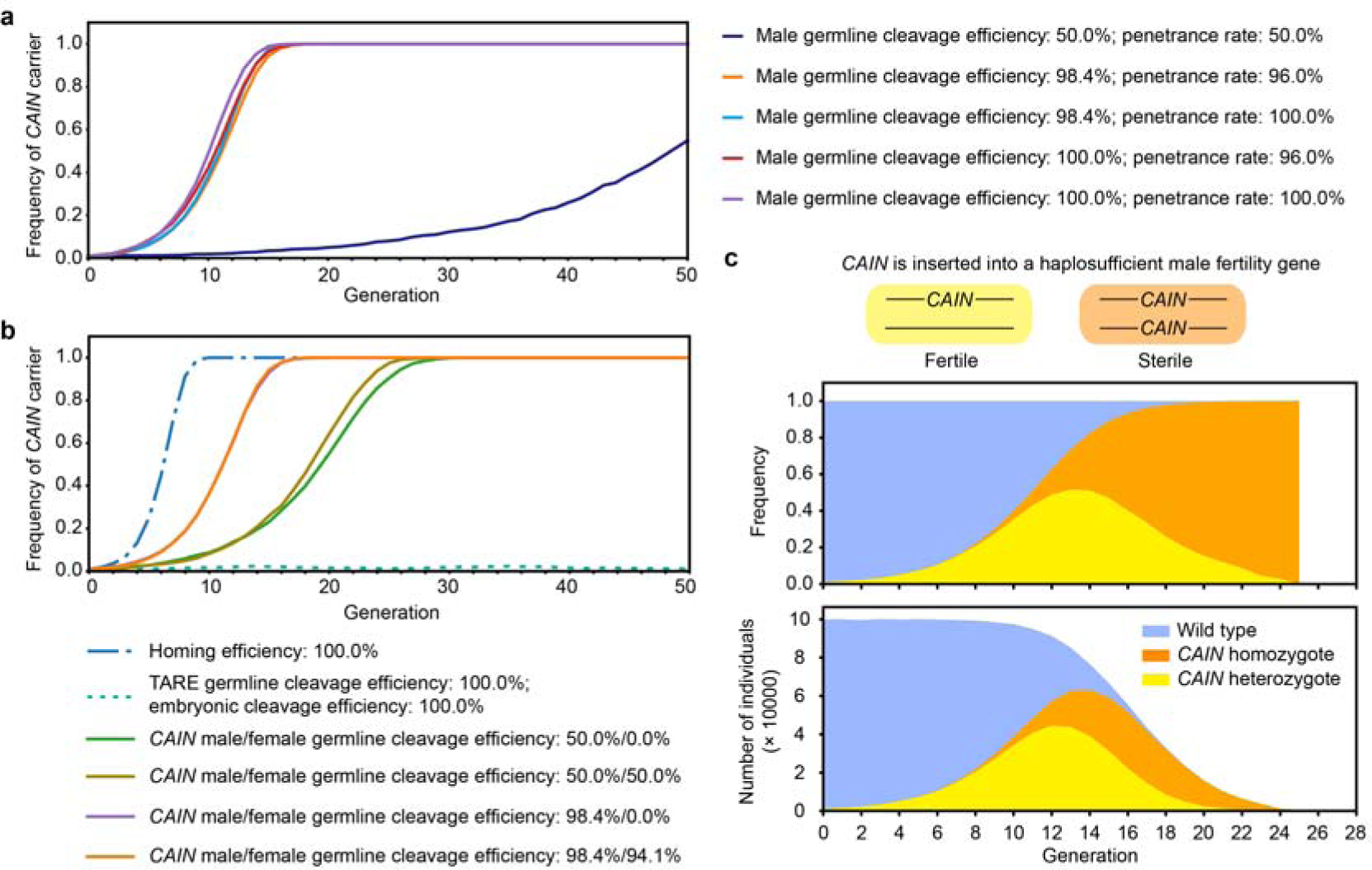
Simulated spread dynamics of modification and suppression *CAIN* drives. **a,** Computational simulation illustrating the impact of different settings for male germline cleavage efficiency (empirical: 98.4%, artificial: 50.0% and 100.0%) and penetrance rate (empirical: 96.0%, artificial: 50.0% and 100.0%) on the spread dynamics of *CAIN*. **b,** Computational simulation of the spread dynamics for the homing-based drive, TARE, and *CAIN*, with an introduction frequency of 1%. The efficiency of the homing-based drive and TARE was set at their maximum values. For *CAIN*, the penetrance rate was set to the empirically estimated value of 96.0%. Various germline cleavage efficiencies were applied for male (empirical value of 98.4% or artificial value of 50.0%) and female (empirical value of 94.1% and artificial values of 0.0% and 50.0%). **c,** The spread dynamics of the suppression *CAIN* drive. The graph shows the changes in the number and frequency of *CAIN* carriers and wild-type individuals across generations.

### Prevalent allelic conversion of *NPG1* in the germline of *TPD-CAIN* carriers

A striking observation was the consistent absence of the *NPG1*^+^ genotype detectable through Sanger sequencing in somatic tissues of FAST+ F1 plants resulting from pollination of a wild-type plant (*NPG1*^+/+^) with *TPD-CAIN*; *NPG1^−^* pollens (Fig. 3b; Table S4, *n* = 49). It suggested that the maternal *NPG1*^+^ allele likely underwent CRISPR-mediated DSB. To further test this hypothesis, we conducted a thorough analysis of *NPG1* genotypes around the gRNA11 target site using Illumina sequencing on leaf and inflorescence samples from three FAST+ F1 plants derived from male parent T18. Besides the primary (averaging 52.9%) genotype “*NPG1*^−8^”, likely inherited from T18 (Fig. 3b), several other genotypes were also observed, although the un-cleaved *NPG1*^+^ allele was sparse (averaging 0.8%, Fig. 3c). This diversity among *NPG1* alleles implied that DSBs in the maternal *NPG1* allele were induced by Cas9 after the one-cell zygote stage, with these DSBs primarily repaired via end-joining mechanisms in somatic cells.

The Cas9 activity responsible for inducing these DSBs could originate from one of the two plausible scenarios: *Cas9* expressed from the embryo’s genome post-fertilization (*i.e.*, zygotic), or the paternal carryover of Cas9 proteins. The latter scenario seemed less likely due to limited protein/RNA content allowed by the small sperm cell size^30^. To investigate these two hypotheses, we examined the *NPG1* genotype of F2 plants generated by the reciprocal F1 backcrosses conducted earlier (Fig. 4), using Sanger sequencing (Fig. 6c; Table S5). The *NPG1*^+^ genotype seldom appeared in FAST+ F2 progeny, regardless of *TPD-CAIN* inheritance from either the male or female parent, occurring only 2/17 times (gRNA2 target site) and 0/17 times (gRNA11 target site) when inherited from male parents, and 0/25 times at both gRNA2 and gRNA11 target sites when inherited from female parents (Table S5). In contrast, the *NPG1*^+^ genotype was detected in all FAST− F2 progeny, even if the male parent carried *TPD-CAIN* (Fig. 6c; Table S5). These observations aligned well with zygotic or later Cas9 expression, contradicting the explanation of paternal carryover.

Analyzing the transmitted *NPG1* genotypes from FAST+ F1 to F2 plants (Fig. 6c; Table S5) revealed a frequent dominance of a single *NPG1* genotype (with a frequency far exceeding 50%), despite these F1 parents being initially heterozygous *NPG1^+/−^* at zygote formation. For instance, although three distinct *NPG1* genotypes were transmitted from the same FAST+ F1 male parent T18-2-1 (*NPG1*^−8^) to F2 plants at the gRNA11 target site, *NPG1*^−8^ appeared in 16 out of 19 F2 plants (Table S5). The remaining two genotypes, *NPG1*^−C^ and *NPG1*^−30^, were identified in only two and one plant, respectively (Table S5). This pattern held true regardless of whether FAST+ F1 plants served as male or female parents, with the primary *NPG1* allele being transmitted to F2 plants at frequencies of 77% and 90%, respectively (Table S5). This skewed representation could not be simply attributed to ending-joining repair bias, as it occurred with two different *NPG1* alleles, *NPG1*^−C^ and *NPG1*^−8^, transmitted from T06 and T18, respectively (Table S5). These findings suggested that, in contrast to somatic tissues, HDR was preferentially used to repair DSBs in germline cells, using the paternal *NPG1*^−^ allele as a template and resulting in allelic conversion.

### Enhancing *CAIN* dissemination through *NPG1* cleavage within the female germline

The process of *NPG1* cleavage and its subsequent repair within female germlines has the potential to enhance the spread of *CAIN* by complementing the *NPG1* knockout achieved in male germlines. To obtain an accurate estimation of *NPG1* cleavage efficiency in female germlines, we analyzed the genotypes of FAST− F2 plants, which provided the genotypic information of the original *NPG1* alleles inherited from their parents, given that no additional *NPG1* cleavage would have occurred post-F2 zygote formation. Among the 34 FAST− F2 progeny resulting from the cross-pollination of FAST+ F1 plants (*i.e.*, *TPD-CAIN*/+) with wild-type pollen, 33 exhibited a heterozygous *NPG1*^+/−^ genotype at the gRNA11 target site (Table S5), higher than the 50% expected in the absence of Cas9 cleavage occurring in female germlines. The prevalence of the *NPG1*^−^ allele, likely introduced through the female gamete, suggested a female germline cleavage efficiency of 94.1% (Fig. S5b). All offspring had paternal wild-type alleles, indicating minimal maternally deposited Cas9 activity, in contrast to several insect CRISPR gene drive systems^17^. While such activity could somewhat benefit *CAIN*, it is not a crucial factor^24^.

To assess how this female germline *NPG1* cleavage could assist in *CAIN* spread, we incorporated this parameter into a simulation with an initial introduction frequency of *CAIN* carriers set at 1% (Fig. 7b). The results revealed a scenario-dependent impact: when male germline cleavage efficiency is at 50%, an additional 50% female germline cleavage could expedite *CAIN* propagation by approximately 3 generations. However, at the observed 98.4% male germline cleavage efficiency in *TPD-CAIN* plants, the actual female germline cleavage efficiency of 94.1% only provided a marginal advancement of roughly one generation to *CAIN* propagation (Fig. 7b). The simulation results also indicated that the rate of *CAIN* spread was only delayed by a few generations compared to invasive, homing-based drives. However, it remained significantly faster than the frequency-dependent TARE drive, which requires moderate release sizes to quickly increase in frequency (Fig. 7b).

*CAIN*/TADS has significantly faster propagation compared to TARE at higher frequency^18^, which underscores its potential as a tool for population suppression when integrated into a haplosufficient male fertility gene that is essential exclusively before the haploid stage^24^. To gauge its effectiveness in suppression, we conducted simulations on the population dynamics of *CAIN* with an initial release frequency of 1% (Fig. 7c). These simulations showed an initial rapid increase in both the number and frequency of *CAIN* carriers, specially heterozygotes. Simultaneously, the total population size decreased progressively, ultimately resulting in population extinction by generation 26 (Fig. 7c), likely due to the surge in male-sterile *CAIN* homozygotes. Collectively, these simulation results indicated that *CAIN*’s rapid propagation could allow it to be configured as a potent suppression drive.

## Discussion

Mendelian inheritance, which is characterized by the equal probability of inheriting either allele from heterozygous diploid parents, promotes efficient adaptation through natural selection^31^. This same mechanism, however, prevents traits that are advantageous to humans yet harmful to the organisms themselves from spreading in wild populations. The *CAIN* synthetic toxin-antidote gene drive system overrides Mendelian inheritance, thereby enabling the propagation of genes favored by humans. It is worth noting that earlier synthetic toxin-antidote gene drives, such as synthetic *Medea*^32,33^ and *ClvR*^17^/TARE^18^ were primarily designed to bias inheritance in female parents, and the resulting embryonic nonviability compromised fertility. In contrast, targeting the male germline can mitigate potential fitness impacts on fecundity because of the higher abundance of pollen grains compared to ovules, facilitating rapid drive propagation even at low frequencies. Nonetheless, there remains a possibility that the overall fertility of drive heterozygous plants might be diminished if their pollen grains potentially fertilize a smaller number of female gametes due to the failure of germination of some male gametes lacking the drive. In this scenario, modification drives would continue to operate effectively even with moderate fertility decline, whereas suppression drives relying on *CAIN* would experience more pronounced challenges^24^.

*CAIN*/TADS gene drives offer several advantages over more extensively studied homing-based drives. Notably, these gene drives can utilize a wider variety of Cas9 promoters while maintaining a higher propagation speed^24^. Moreover, their cargo genes are likely to possess greater genetic stability due to the potentially lower occurrence of DNA replication errors compared to the copy-and-paste HDR mechanism employed by homing-based drives. Additionally, a suppression drive of the *CAIN*/TADS type would likely exhibit significantly enhanced performance compared to homing-based drives and other commonly considered suppression mechanisms like X-shredders^34^ when spatial structure is taken into account. This enhancement is manifested both in terms of ease of achieving a high level of suppressive power and in preventing the persistence of wild-type alleles^35^.

Using the FAST marker^29^ as the cargo gene, we established *CAIN* as a state-of-the-art tool to efficiently modify entire plant populations. The cargo is versatile, adaptable for various ecological challenges, with gene selection dependent on specific scenarios and goals (Fig. S6). For instance, *CAIN* can boost inheritance of advantageous traits—such as drought or disease resistance genes^36,37^—to strengthen the resilience and survivability of endangered, fragile plant species in their natural habitats, potentially heralding a new era in ecological management. Furthermore, the rise of herbicide-resistant weeds often prompts increased herbicide usage^38,39^, which in turn threatens biodiversity and potentially reduces productivity through broad spectrum effects. By incorporating genes that counter herbicide resistance or by disrupting herbicide resistance genes, the effectiveness of a weed control can be maintained while reducing the need for excessive spraying, regardless of emerging resistance. This gene drive-based approach thus seeks to balance crop protection and environmental considerations to minimize the loss of biodiversity while optimizing productivity.

Acknowledging the ongoing intense debate and regulatory oversight of gene drive technologies^6,40-42^, we prioritized biosafety in the development of *CAIN*. *A. thaliana* was selected as our self-pollinating model plant to prevent the inadvertent spread of *CAIN*, as gene drives require outcrossing to effectively spread throughout a population. The design of the *CAIN* system allows high specificity by enabling the targeting of distinct genotypes or ecotypes, achieved by selecting gRNA target sites based on naturally occurring genetic polymorphisms. The *CAIN* system is intrinsically zero-threshold, meaning it can spread throughout a population with just a few drive-carrying individuals, making it less confined compared to other CRISPR toxin-antidote drives like TARE. Nevertheless, methods proposed to limit homing drive spread^43^ can be integrated with *CAIN*. For example, by propagating a confined drive (*e.g.*, TARE) to supply the required Cas9, the *CAIN* drive can be “tethered” to a specific geographic area by a TARE drive. Such flexibility allows for a measured application of *CAIN*, offering tunable levels of confinement for different scenarios and thus enabling secure application across diverse ecological contexts.

Updatability is a critical safety aspect for gene drives. The design of *CAIN* incorporates the capacity for functional replacement, providing a means to modify or remove existing gene drives, as necessary. This process, outlined in Fig. S7, depends on three prerequisites. First, the successor *CAIN* drive (*CAIN_n_*_+1_) should integrate at the same genomic locus as its predecessor (*CAIN_n_*), placing the two in direct competition. Despite the low rate of targeted integration via HDR^44^ in plants, this problem can be offset by screening a large transformant population. Second, *CAIN_n_*_+1_ uses distinct gRNA target sites to disrupt the essential gene, *NPG1*, thus acting as a next-generation toxin that can target both the recoded essential gene in *CAIN_n_* as well as the wild-type *NPG1* (though targeting the wild-type may not be necessary if the original drive has disrupted all wild-type alleles). Third, an updated recoded essential gene that is resistant to the current and next-generation CRISPR cleavage can act as the next-generation antidote. Once all these conditions are in place, a *CAIN_n_*_+1_ drive can be used to override *CAIN_n_*, enabling changes or removal of cargo genes. These advantages also preclude the need for a new target gene, preserve the size of the gene drive construct, and may ultimately boost integration efficiency. A similar strategy has been experimentally demonstrated for the *ClvR* system^45^.

In summary, we introduce *CAIN*, a CRISPR-based toxin-antidote gene drive system tailored for plant species. This innovation leverages the distinct features of plant biology, particularly the extended haploid stage in the plant life cycle, to potentially shift the genetic composition of wild plant populations. Our successful demonstration of *CAIN* in *A. thaliana* sets a precedent for its application to other plant species, since *NPG1*, a critical component of our system, is highly conserved across a wide range of monocots and dicots^46^. Looking forward, further refinement of the *CAIN* gene drive is imperative, including the development of strategies for its reversibility and adaptability, as well as exploring possible mechanisms for its genotypic or geographical confinement. As we venture into this new frontier in genetic engineering, *CAIN* and similar gene drive systems could reshape ecological management, agricultural practices, and initiatives for the preservation of endangered plant species.

## Methods

### Materials and growth conditions

All *A. thaliana* lines used in this study were derived from the Columbia-0 (Col-0) background. Seeds were surface sterilized with 10% sodium hypochlorite for 10 min, followed by three rinses with sterile water, before being sown on germination medium (2.2 g/L^−1^ Murashige and Skoog medium, 10 g/L^−1^ sucrose, and 7.6 g/L^−1^ plant agar, pH = 5.7). After stratification at 4°C for 2 days, the agar plates were transferred to a growth chamber set at 22°C under a 16-h light/8-h dark photoperiod. Seven-day-old seedlings were then transplanted into soil and maintained in the greenhouse under the aforementioned conditions for further experiments.

### Construction of plasmids

All restriction enzymes and enzymes for Gibson assembly were obtained from New England Biolabs (NEB). Hi-fidelity polymerase (Phanta Max Super-Fidelity DNA Polymerase, P505) for polymerase chain reaction (PCR) and gel extraction kit (FastPure Gel DNA Extraction Mini Kit, DC301) were obtained from Vazyme. All the plasmids used in this study were cloned using standard molecular biology techniques and purified using StarPrep Fast Plasmid Mini Kit (Genstar, D201). All primers used in this study are listed in Table S6.

The construction of the FAST-only vector involved inserting the *FAST* marker^29^ (*pOLE1*:*OLE1*-*TagRFP*-*Nos terminator*) into the XF675 binary vector through Gibson assembly^47^. To achieve this, the promoter and partial coding sequences of the *OLE1* gene (AT4G25140) were amplified from genomic DNA (gDNA), while *TagRFP* and *Nos terminator* sequences were synthesized as provided in a previous study^29^. The HindIII-digested XF675 plasmid, along with the two fragments mentioned above, each having 20 bp overlaps with adjacent fragments, were assembled using Gibson assembly.

To generate *DMC-CAIN* and *TPD-CAIN* vectors containing *pDMC1*/*pTPD1*:*Cas9*-4_gRNA_cassette-FAST-Rescue (*i.e.*, *pNPG1*:*recoded NPG1*), all the components were cloned into the XF675 binary vector using Gibson assembly or golden gate ligation in five successive steps. First, the sgRNA cassette (*pU6-SmR-gRNA scaffold-U6 terminator*) was amplified from the plasmid pHEE401E (Addgene plasmid #71287) and inserted into the double-digested XF675 binary vector (digested with EcoRI and HindIII) using Gibson assembly. The HindIII recognition site was made complete for following digestion. Second, four gRNA cassettes, each driven by a U6 promoter (U6-26, U6-29, U6-1, and U6-26), were multiplexed using BsaI digestion and golden gate ligation as described^28^. Third, the resulting plasmid was then digested with HindIII, and the FAST marker sequences were cloned in using Gibson assembly, with the HindIII recognition site completed again. Fourth, the promoter (2427 bp, amplified from gDNA) of *NPG1*, the recoded coding sequence (without introns) obtained by PCR mutagenesis, the native 3’ UTR (166 bp, amplified from cDNA) and the NOS terminator (amplified from the plasmid XF675) were subsequently inserted into the HindIII-digested plasmid using Gibson assembly. Fifth, the resulting plasmid was digested with KpnI, and the following fragments were cloned in using Gibson assembly: the respective promoter sequences of *DMC1* (AT3G22880, 2172 bp) and *TPD1* (AT4G24972, 2770 bp) amplified from gDNA, the plant-codon-optimized *SpCas9* sequence amplified from the plasmid pHEE401E, and the NOS terminator sequences amplified from the plasmid XF675.

### Design principles for *CAIN*

Implementing a successful toxin in *CAIN* requires a gene to be essential for pollen germination. To identify suitable target genes, we retrieved candidates from a previously compiled list of genes important for male gametophyte development, but which do not affect female gametophyte development^48^. *No Pollen Germination 1* (*NPG1* or AT2G43040), which is essential for pollen germination^25^, was selected as the target for our gRNA-Cas9 cassettes because its later role in male gametophyte functionality should allow more time for CRISPR-based gene knockout to occur than genes with earlier stage functions (Fig. 1a). The antidote was designed using a recoded version of *NPG1*, driven by its native promoter, to restore normal pollen germination. To ensure resistance to further Cas9 cleavage, we introduced 28 synonymous mutations around the four gRNA target sites (Fig. S1a). Additionally, introns were removed from the recoded version to ensure that *CAIN* constructs were as small as possible (Fig. S1b).

### Evaluating the editing efficiency of potential target sites in protoplasts

The CRISPR-P 2.0^49^ bioinformatics tool (http://crispr.hzau.edu.cn/CRISPR2/) identified 35 potential Cas9-mediated DNA cleavage target sites within the *NPG1* coding sequence. These were further refined based on four criteria: 1) location within the exon region; 2) positioning within the upper two-thirds of the coding sequence; 3) having fewer than four consecutive thymine bases within the gRNA sequence; and 4) lacking more than 7-base pairing with the scaffold sequence, predicted by CRISPR-GE^50^ (http://skl.scau.edu.cn/home/). To evaluate the editing efficiencies of the remaining 12 gRNA target sites in *A. thaliana* protoplasts (Table S1), the 20-nt gRNA spacer sequences were individually incorporated into the *pAtU6-sgRNA* vector (Addgene plasmid #119775) using BsaI digestion and golden gate ligation.

*A. thaliana* protoplasts were prepared following procedures described in previous studies^51,52^. After co-transformation with *pAtU6-sgRNA* and *p2X35S-Cas9*, protoplasts were cultivated at room temperature for 48 h before harvest. In parallel, another set of protoplasts was transformed with *p2X35S-GFP* alone to measure the transformation efficiency approximately 16 h post-transformation. The transformation efficiency was found to be 41% and 45% in two biological replicates, respectively.

Genomic DNA was extracted from each tube of protoplasts using Plant Genomic DNA Kit (DP305, TIANGEN). The genomic regions spanning 180–220 bp around each target site were PCR amplified and purified. To differentiate PCR products from two biological replicates at the same target site, a 6-nt barcode (ATGCAG) was introduced for one of the replicates using primers (Table S6). Upon purification, each of the 24 PCR product samples was quantified using Nanodrop 2000, pooled together in equal amounts, and then subjected to library construction and Illumina paired-end 150 bp (PE150) sequencing (NovaSeq 6000).

A total of 7,134,194 sequencing reads were obtained from the 24 PCR products. After filtered using fastp^53^ (version 0.23.1) with the parameter “fastp -g -q 5 -u 50 -n 15 -l 150 --min_trim_length 10 --overlap_diff_limit 1 --overlap_diff_percent_limit 10”, 7,055,416 clean reads were retained. The data for the two biological replicates were partitioned according to the presence or absence of the barcode sequence “ATGCAG”, and then mapped to *NPG1* reference sequence using Burrow-Wheeler Aligner^54^ (version 0.7.17-r1188) with the parameter “bwa mem -M”. Variants were called from the BAM files using SAMtools^55^ (version 1.13) with the parameters “samtools mpileup -A -d 0 -Q 13 -q 30 -B”. The depth of reads covering the twelve gRNA regions spanned from 57,412–163,671 in replicate 1 and from 133,643–203,234 in replicate 2, respectively. The mpileup files were then transformed into VCF files using an in-house perl script (https://github.com/QianLabWebsite/GeneDrive).

To identify CRISPR-mediated editing events, we considered variants exhibiting mismatches in the 23-nt spacer and PAM region. While the mismatches could potentially arise from Illumina sequencing errors, most identified mismatches were indels. In contrast, Illumina sequencing errors usually result in single base substitutions (10^−3^ mismatches per base^56^) rather than indels (10^−6^ mismatches per base^57^). To further mitigate errors in variant identification, a frequency threshold of 0.5% was set of single base substitutions and all indels were identified as edited variants. The editing efficiency for each gRNA was estimated by calculating the ratio of mismatched reads to the total number of reads mapped to the respective target site (Table S1).

### Generation of single-locus T-DNA inserted transformants

Wild-type Col-0 plants were subjected to the floral dipping transformation^58^ using *Agrobacterium* GV3101 strain containing one of the three vectors (*DMC-CAIN*, *TPD-CAIN*, or *FAST only*). Successful primary transformants (T1) were selected based on the presence of red fluorescence in dry seeds, detected by a handheld fluorescence detector (LUYOR 3415RG).

To identify single-locus T-DNA inserted transformants, thermal asymmetric interlaced (TAIL-)PCR^59^ was conducted on 48–50 randomly chosen T1 plants from each drive construct. Further validation was carried out using whole genome sequencing (∼20X coverage, NovaSeq 6000 PE150, Novogene) on candidate T1 lines. We developed an in-house pipeline for detecting T-DNA insertion sites. We first created a reference file by concatenating T-DNA sequence with the *A. thaliana* reference genome^60^ (TAIR 10). Sequencing reads were then aligned to this reference using BWA with parameters “bwa mem -M”. We examined chimeric reads (*i.e.*, reads with subsections aligning to separate positions in the reference genome) and discordant read-pairs (pairs with unexpected distance/orientation in their alignments) mapped to the T-DNA borders (300 bp). The residual sequences in these reads (or read pairs) helped us pinpoint putative insertion sites in the genome. We performed local assembly of reads around each putative insertion site using Megahit^61^ (version 1.2.9) with parameters “megahit --prune-level 1 --prune-depth 1 --low-local-ratio 0.1”, and the resulting contigs were aligned to the integrated reference with Blastn^62^ (version 2.9.0+). This allowed us to reconstruct the junction between T-DNA and the chromosome at the insertion site. Finally, we confirmed single-locus T-DNA insertion in three *DMC-CAIN*, four *TPD-CAIN*, and two *FAST only* T1 lines (Table S2).

### Cross-pollination and the estimation of transmission rate

We assessed the transmission rate of the *CAIN* gene drives and the *FAST-only* construct through cross-pollination experiments, quantifying the percentage of progeny seeds that displayed red fluorescence. For cross-pollination, we transferred pollen from 2–3 male parent flowers to the stigma of an unopened, emasculated female parent flower.

*R* (version 4.2.3) and RStudio software (version 2023.03.0+386) were used for statistical analyses. We assessed deviations from Mendelian inheritance by examining the fraction of FAST+ seeds through two-sided exact binomial test, setting a null hypothesis of 50%. Considering potential heterogeneity in *NPG1* cleavage status across male parent flowers, and consequent variations in the proportion of FAST+ seeds across individual siliques, we employed a replicated *G*-test from the DescTools package to detect distortion in the transmission ratio. To account for an increased risk of type I error (false-positive) due to multiple comparisons, we calculated the false discovery rate (FDR) using the p.adjust() function.

### Genotyping

The genotypes at the *NPG1* locus were determined by Sanger sequencing of a ∼2 kb PCR product, amplified from genomic DNA and covering all four gRNA target sites (Fig. S1b; primers detailed in Table S6). DNA was extracted from rosette leaves and early inflorescence (with flowers unopened). Consistent Sanger sequencing results were usually obtained from leaf and inflorescence samples of the same plant. However, if multiple peaks were evident in the chromatograph of leaf samples (indicating potential mosaicism or heterogeneity), genotypes were determined using inflorescence sample from the same plant. Genotyping of FAST− F1 and F2 plants was performed using only leaf samples; samples with indeterminable genotypes were excluded.

For three F1 plants, we determined the gRNA11 target site genotypes using Illumina sequencing on both leaf and inflorescence samples. PCR products from different tissues, each with a unique barcode introduced during PCR (Table S6), were mixed in equal quantities and sequenced with Illumina PE150. Clean reads, sorted by barcode, were mapped to the genomic sequence around the gRNA11 target site using BWA. The depth of reads covering the gRNA11 region varied between 107,207 and 150,422. We considered variants with mismatches in the 23-nt spacer and PAM region, and determined the editing efficiency per sample as the ratio of reads with mismatches to total reads mapped to the target site. We identified single base substitutions with a frequency above 0.5% and all indels as edited variants.

CRISPR cleavage potential beyond *NPG1*, caused by the incorporated gRNAs in *CAIN*, raises concerns about off-target effects, which could unintentionally introduce genomic mutations and increase the genetic load to the population. We predicted 16 potential off-target sites using CRISPR-P 2.0^49^ and tested the genotypes at these locations in 16 FAST+ F1 plants (*TPD-CAIN*/+; *NPG1*^−/−^) using Sanger sequencing (Table S7). The absence of mutations at these potential off-target sites in all the tested plants indicated that the four gRNAs in *CAIN* specifically targeted *NPG1* as designed.

### Simulation of the population dynamics

The population dynamics of *CAIN* were computationally simulated using an individual-based stochastic model based on the Wright-Fisher model, which assumes a constant population size and non-overlapping generations. For the modification *CAIN* drive, we considered genotypes at two unlinked loci, *CAIN* and *NPG1*, and initiated the population with 9900 wild-type individuals and 100 *CAIN-*carrying heterozygotes (*TPD-CAIN*/+; *NPG1*^+/+^). We implemented a density regulation production strategy following a previous study^24^. Briefly, we calculated a scaling factor (*S*) using the formula *S* = 10 / (9 × *N* / *K* + 1), where *N* represented the population size of the current generation, and *K* represented the environmental carrying capacity (*i.e.*, 10,000). We calculated the population size for each generation (*X*) from a binomial distribution *X* ∼ *B* (50, 0.02 × *S*). For each generation, a pair of individuals were randomly chosen to produce offspring, with the roles of male and female randomly assigned. This process was repeated *X* times, and the resulting offspring replaced the parent generation.

For each generation, male parents carrying *CAIN* could trigger CRISPR-mediated cleavage at the *NPG1* locus, at a rate either estimated from empirical data (98.4%, Fig. S5a) or set artificially (50% or 100%). This cleavage led to the creation of loss-of-function *NPG1* alleles, and the corresponding lethal phenotype in male gametes had a penetrance rate either empirically determined (96.0%, Fig. S5a) or artificially set (100%). We also considered germline cleavage efficiency in *TPD-CAIN*/+ females, set either based on empirical data (94.1%, Fig. S5b) or artificially (0%, 50%, 100%).

For the homing-based drive, the presence of the drive allele in a heterozygous state led to the conversion of the wild-type allele into the drive allele with a 100% efficiency. For the TARE drive, the target gene was a haplosufficient essential gene for embryo development, and Cas9 would cleave the wild-type target gene in the germline with a 100% probability. Following fertilization, maternally carried Cas9 would further cleave the paternal wild-type target gene, also with a 100% probability (*i.e.*, embryonic cleavage efficiency). Embryos bearing two disrupted alleles of the target gene yet without the TARE drive would not proceed to develop.

For the suppression *CAIN* drive, *CAIN* was placed within and consequently disrupted a haplosufficient male fertility gene. The simulation was otherwise similar to that of the modification *CAIN* drive, with the exception that the initial population size and *K* were both set to 100,000, and that the male homozygous *CAIN* plants were incapable of producing viable pollen.

The simulation was implemented using a custom Python script, available at https://github.com/QianLabWebsite/GeneDrive.

## Data availability

The Illumina sequencing data have been deposited in the Genome Sequence Archive^63^ in National Genomics Data Center^64^, China National Center for Bioinformation/Beijing Institute of Genomics, Chinese Academy of Sciences under accession number CRA011573 (temporary link for reviewers https://ngdc.cncb.ac.cn/gsa/s/Tby9C5yR). Plasmid sequences have been deposited in GenBase (accession ID: C_AA 031173.1, C_AA031174.1, and C_AA031175.1).

## Code availability

The scripts developed for variant calling at gRNA target sites and simulation of the population dynamics for *CAIN* are available at Github (https://github.com/QianLabWebsite/GeneDrive).

## Acknowledgement

We thank Dr. Jianzhi Zhang from University of Michigan, Paul Thomas from University of Adelaide, and Dr. Caixia Gao, Dr. Taolan Zhao, and Dr. Yanming Chen from Institute of Genetics and Developmental Biology CAS for discussion. We thank Dr. Xiaofeng Cao from Institute of Genetics and Developmental Biology CAS for providing plasmid XF675.

## Author contributions

W.Q. and Y.L. designed the study; Y.L. performed experiments; B.J. performed computational analyses; Y.L., J.C., and W.Q. wrote the manuscript.

