## Supplementary Figures for "Overriding Mendelian inheritance in *Arabidopsis* with a CRISPR toxin-antidote gene drive that impairs pollen germination"

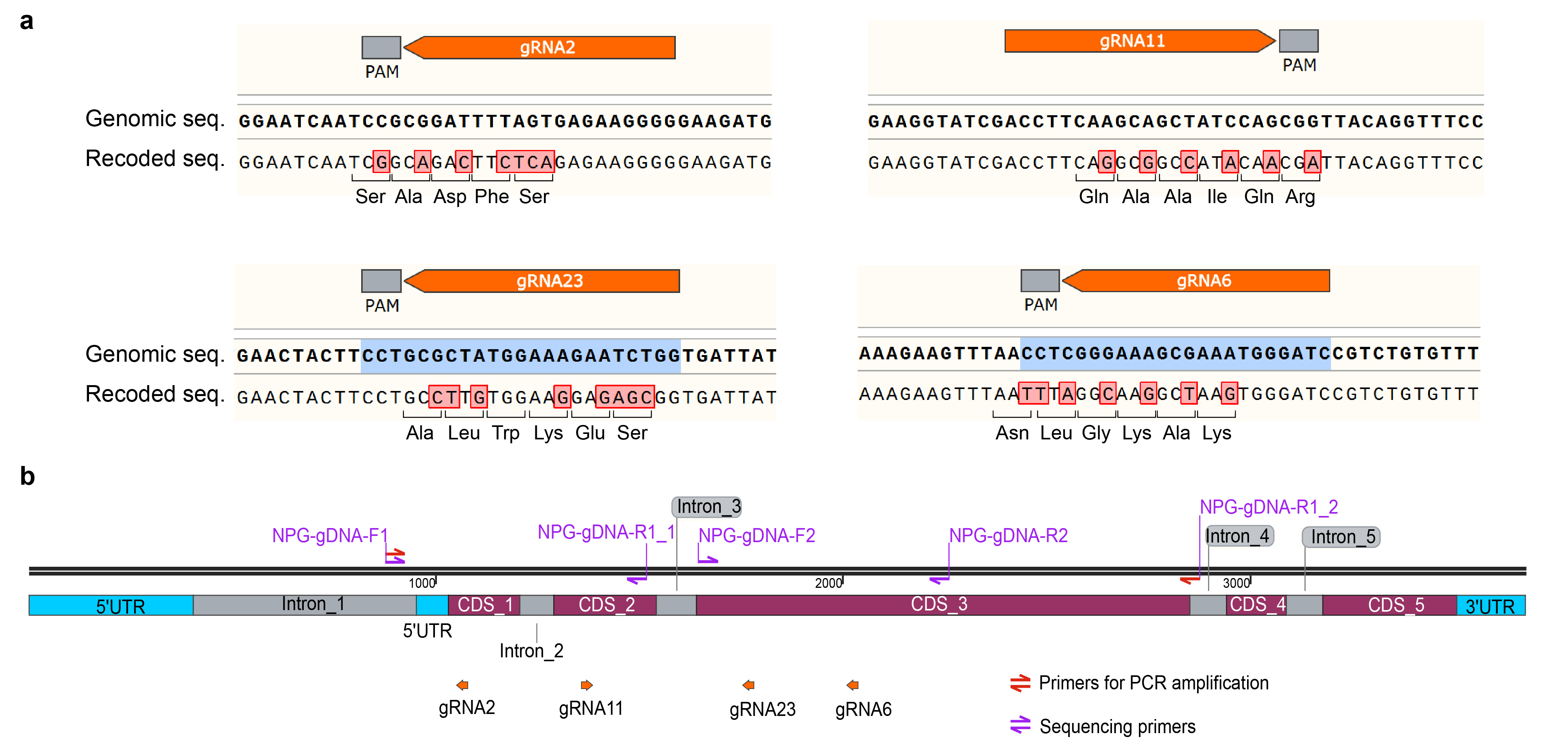

**Figure S1. Incorporation of four gRNAs into *CAIN*.**

**a,** Four gRNAs selected to be multiplexed into the *CAIN* constructs. Mutated nucleotides in the recoded version of *NPG1*, based on synonymous codons, are indicated by red squares. **b,** Genomic positions of the gRNAs and genotyping primers. Amplification of genomic regions containing the four target sites was performed using the NPG-gDNA-F1 and NPG-gDNA-R1_2 primers (in red). Sanger sequencing of the target sites was performed using four primers (NPG-gDNA-F1, NPG-gDNA-F2, NPG-gDNA-R1_1, and NPG-gDNA-R2, in purple).

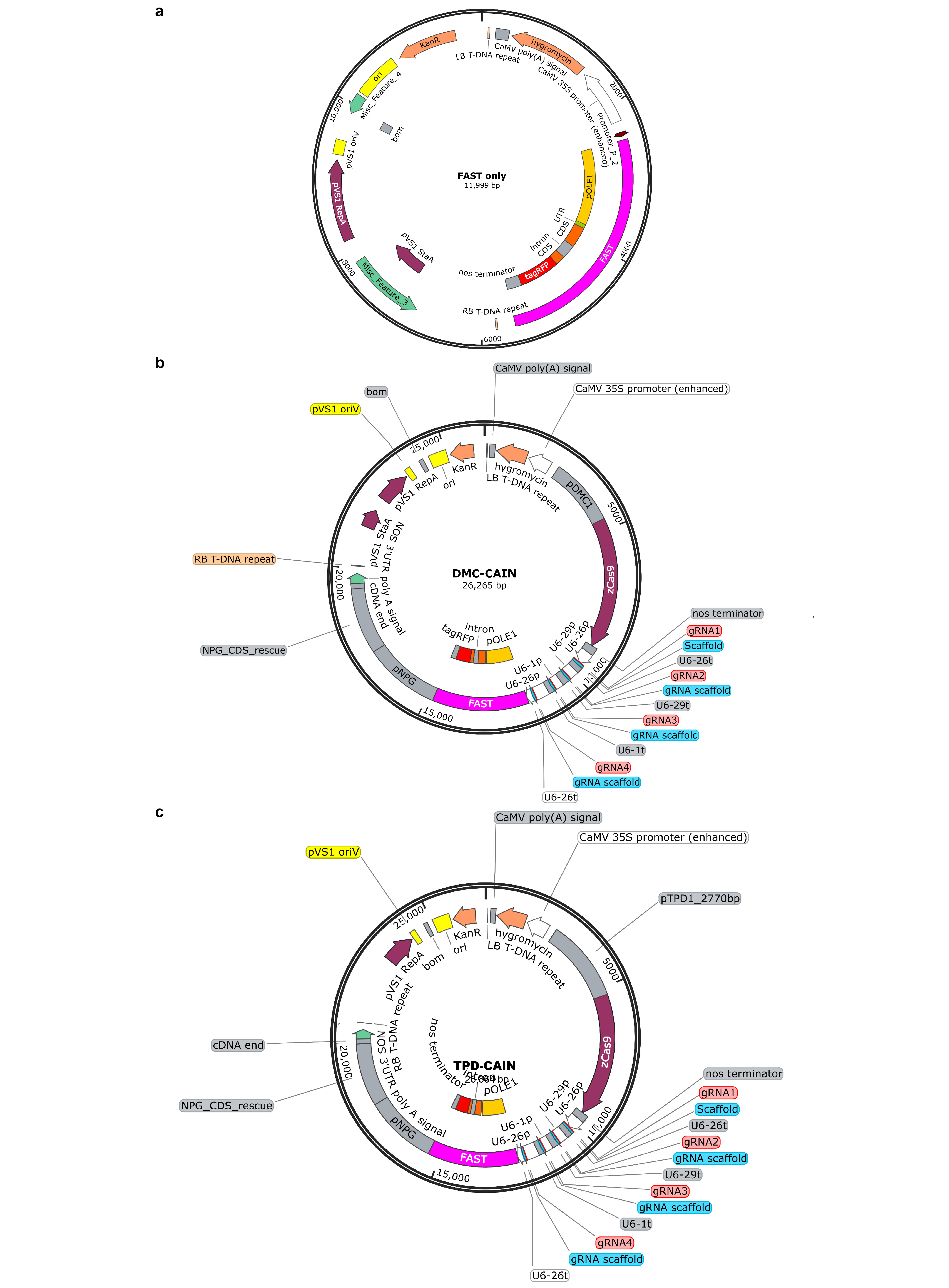

**Figure S2. Schematic maps of *FAST only* control cosntruct and *CAIN* gene drives.**

Shown are schematic maps of the (**a**) *FAST only* control construct, (**b**) *DMC-CAIN*, and (**c**) *TPD-CAIN* gene drive constructs, illustrating their total length and main features.

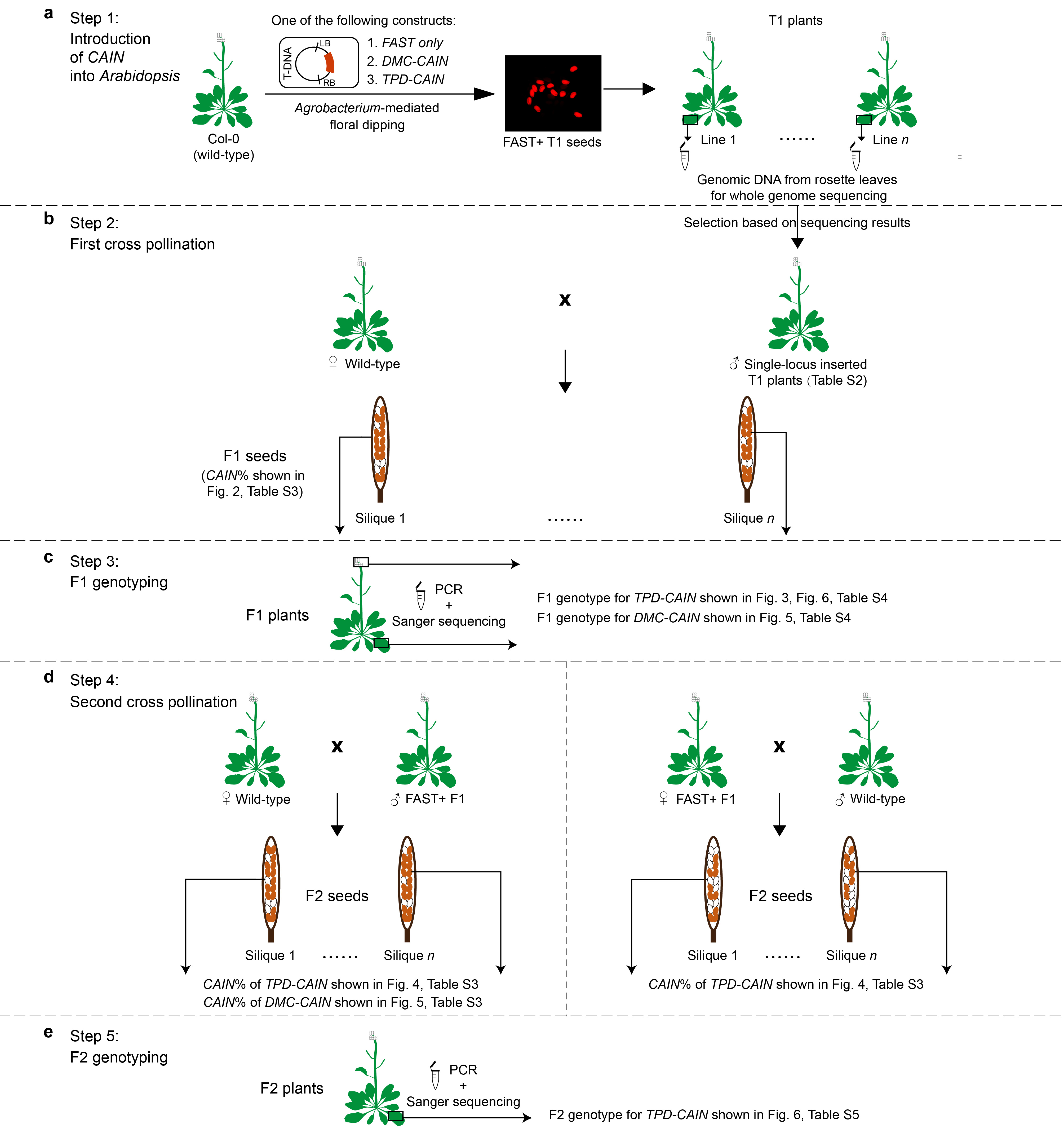

**Figure S3. Experimental procedures overview.**

**a,** Step 1: Introduction of control construct (*FAST only*), *DMC-CAIN*, or *TPD-CAIN* gene drive construct into *Arabidopsis* Col-0 plants, indicated by red fluorescence marker (FAST+) in dry seeds. Single-locus insertion screening was performed using TAIL-PCR and whole genome resequencing. **b,** Step 2: Screened T1 plants were used as the male parent to cross with wild-type female parent. Transmission rate of the *CAIN* gene drive (*CAIN*%) was calculated from the fraction of F1 seeds exhibiting FAST marker. **c,** Step 3: Cultivation of F1 seeds to obtain F1 plants and genotyping (as described in Methods). **d,** Step 4: Crosses between F1 plants of known genotypes (male or female parent) and wild-type plants, calculating *CAIN*% in F2 seeds. **e,** Step 5: Cultivation of F2 seeds to obtain F2 plants and genotyping.

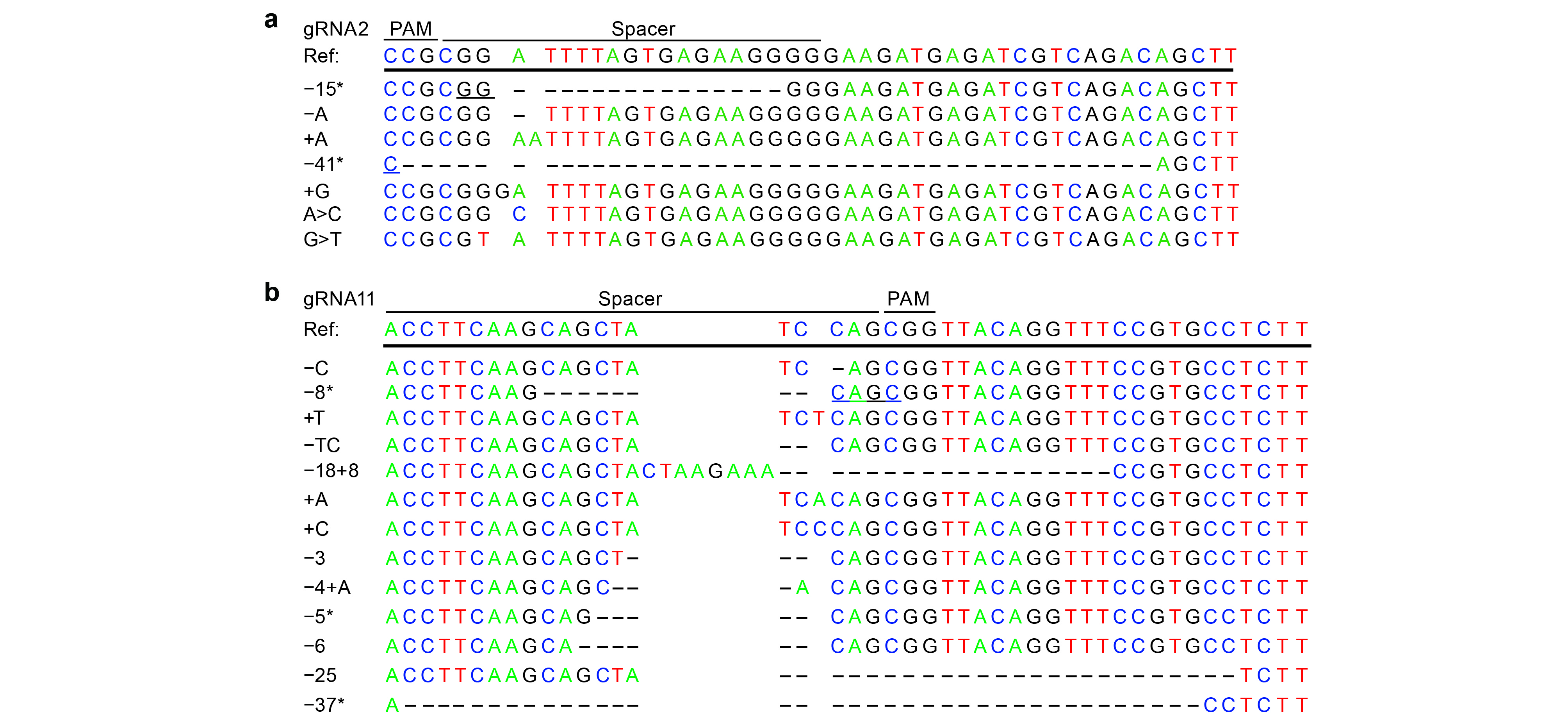

**Figure S4. Mutation types deteced in FAST+ F1 plants.**

This figure illustrates the locations of specific bases invloved in insertions, deletions, and single nucleotide polymorphisms generated at the (**a**) gRNA2 and (**b**) gRNA11 target sites. * denotes one potential alignment result, as the underlined bases can be positioned on either side of the deletion.

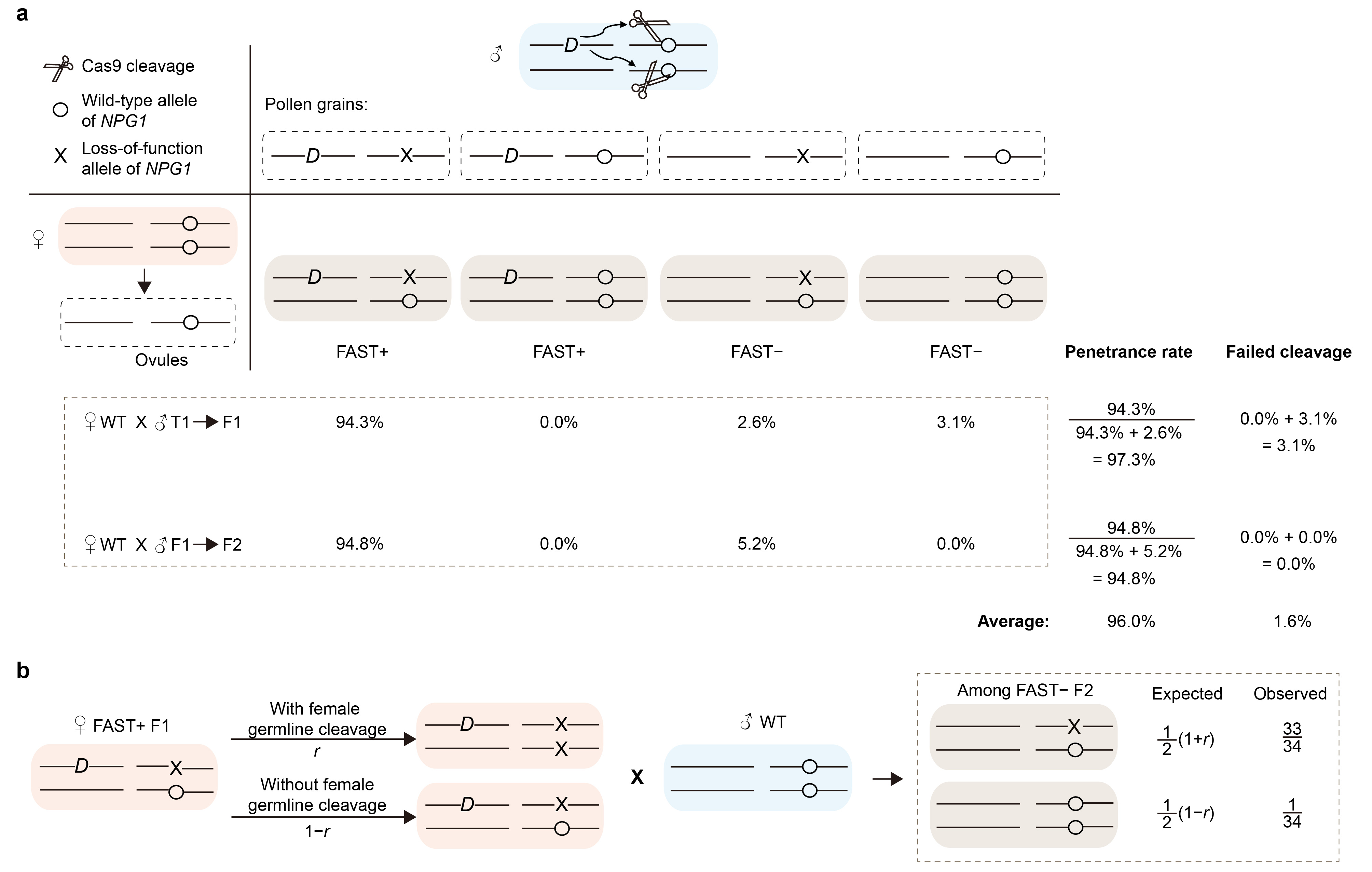

**Figure S5. The impacts of male germline cleavage efficiency, incomplete penetrance, and female germline cleavage efficiency on the *TPD-CAIN* system.**

**a,** Estimated male germline cleavage efficiency and penetrance rate in F1 and F2 progeny. Among the F1 progeny, 94.3% (526/558, Table S3) of the plants were FAST+ (*TPD-CAIN*/+), with all of them showing an *NPG1*^−^ genotype at the gRNA11 target site (Table S4). A small percentage, 2.6% (5.7% × 5 / 11), were +/+; *NPG1*^+/−^ plants, and another 3.1% (5.7% × 6 / 11) were +/+; *NPG1*^+/+^ plants. When examining our F2 progeny, 94.8% (3868 out of 4080, Table S3) were FAST+ (*TPD-CAIN*/+), all of which showed an *NPG1*^−^ genotype at the gRNA11 target site (Table S5). The remaining 5.2% F2 plants were all +/+; *NPG1*^+/−^ (Table S5). From these results, we estimated an average failed cleavage rate of 1.6%, which corresponds to a male germline cleavage efficiency of 98.4%. Additionally, we estimated an average penetrance rate of 96.0%. **b,** Estimation of female germline cleavage efficiency (*r*). We calculated the cleavage efficiency *r* using the genotypes of the *NPG1* locus at the gRNA11 target site in FAST− F2 plants, assuming no further cleavage occurs in these plants. From an observed fraction of 1/34 +/+; *NPG1*^+/+^ plants (Table S5), we estimated *r* to be 94.1%.

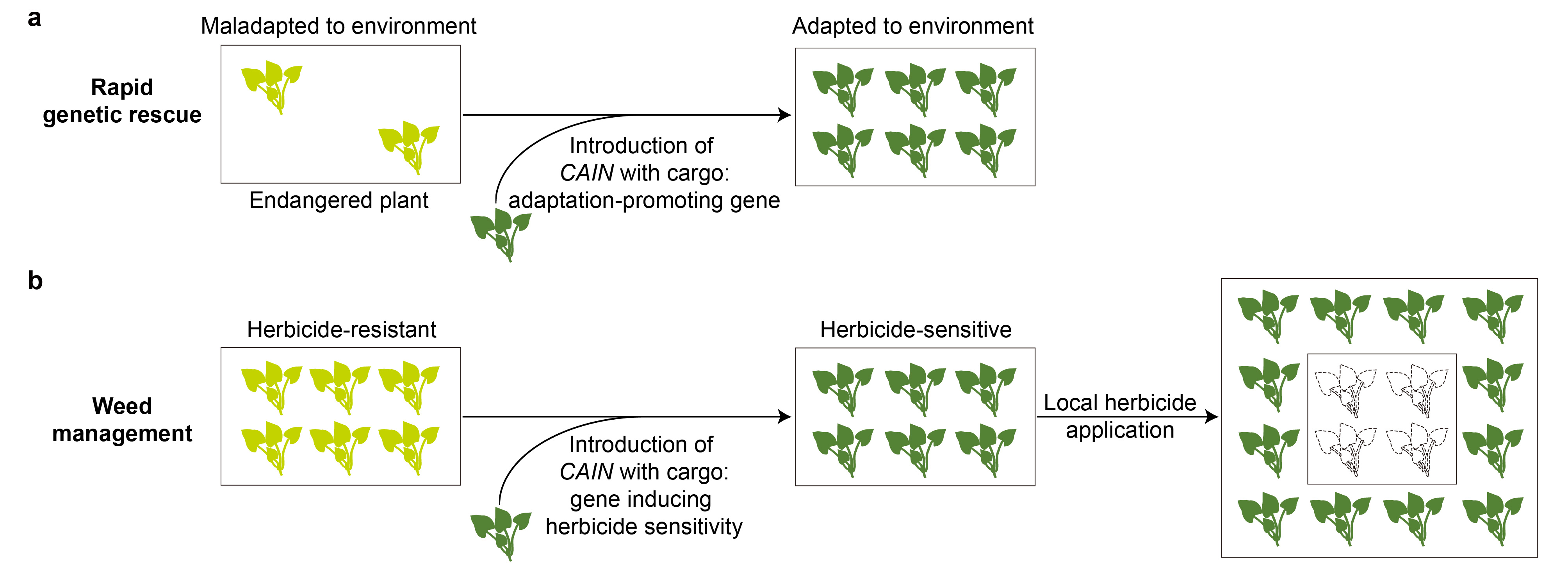

**Figure S6. Potential applications of *TPD-CAIN* gene drive.**

*TPD-CAIN* offers two potential applications: **a,** Adaptation promotion: by incorporating cargo genes that promote the adaptation of specific endangered species, *TPD-CAIN* facilitates rapid genetic rescue and enables the targeted species to adapt to their environment. **b,** Weed management: by incorporating cargo genes that confer herbicide sensitivity to targeted weeds, *TPD-CAIN* allows localized herbicide application for regional weed suppression, providing an additional level of control for gene drive spread.

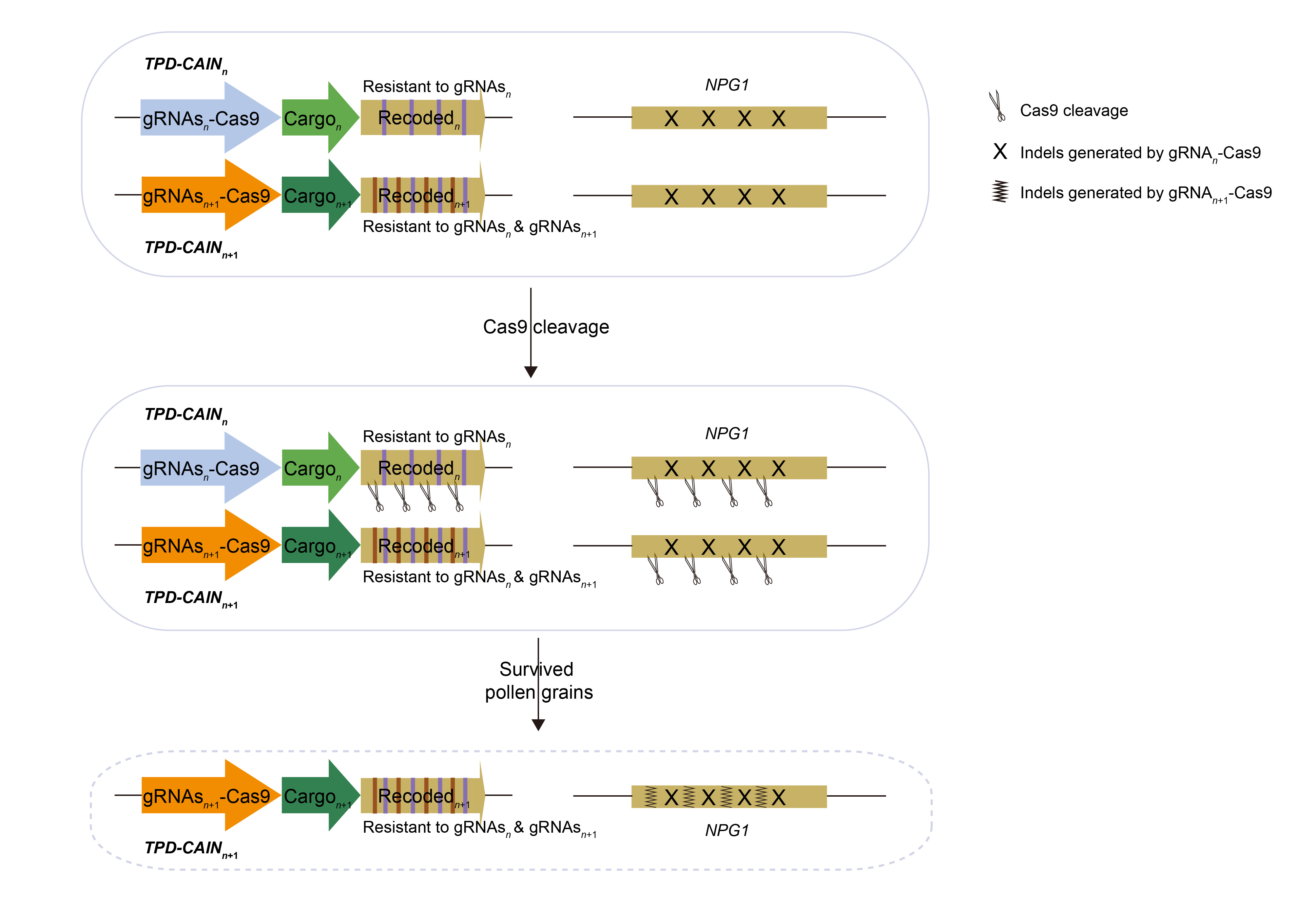

**Figure S7. Updatability of *TPD-CAIN* gene drive.**

In the next generation gene drive, *TPD-CAIN_n_*_+1_, new gRNAs*_n_*_+1_ targeting additional sites within *NPG1* serve as the new toxin, while recoded*_n_*_+1_ that are resistant to both gRNAs*_n_* and gRNAs*_n_*_+1_ serve as the antidote. During male germline cleavage, both the genomic loci and recoded*_n_* would be disrupted by the new toxin, necessitating the rescued activity provided by recoded*_n_*_+1_. This process compels the replacement of the previous generation *TPD-CAIN_n_* with *TPD-CAIN_n_*_+1_, facilitating the spread of the new cargo*_n_*_+1_ linked to the gene drive.
